# Genomic plasticity drives olfactory adaptation in a pest fly

**DOI:** 10.1101/2025.09.18.677102

**Authors:** Qi Xue, Hany K. M. Dweck

## Abstract

Preference shifts in insects are often driven by changes in the olfactory system, yet the underlying mechanisms remain unclear. The worldwide pest *Drosophila suzukii*, which oviposits in ripe rather than overripe fruits, provides a powerful model to study these mechanisms and their behavioral consequences. Here, we show that this shift is linked to functional remodeling in four olfactory receptor neurons: ab2B, ab3A, ab4B, and ab10A. While ab3A and ab10A exhibit tuning changes shared with the non-pest relative *D. biarmipes*, ab2B and ab4B display species-specific adaptations in *D. suzukii*. These changes result not only from receptor sequence divergence but also from novel innovations: receptor co-expression in ab3A and partitioned expression of Or67a paralogs in ab2B and ab10A. Together, these findings show how genomic plasticity in chemosensory gene families enables rapid sensory adaptation and niche transition.

## INTRODUCTION

The olfactory system mediates the interactions between insects and their environment. It guides critical behaviors such as foraging, mate choice, and oviposition ^1–4^. These behaviors are tightly linked to survival and reproductive success. This makes the olfactory system a key target of natural selection. The olfactory system, as a result, evolves rapidly in response to ecological shifts, particularly those involving changes in habitat, food resources, or reproductive sites ^5–7^. However, the molecular mechanisms underlying olfactory adaptations remain poorly understood. While numerous studies have documented the remarkable variation in the copy number of chemosensory receptor genes, driven by gene gains and losses, across insect taxa^8–12^, fewer have connected this genomic diversity to functional differences at the receptor and neuronal levels, or to shifts in behavior ^6^.

The worldwide fruit pest *D. suzukii* presents a powerful model to explore these evolutionary links. Unlike most *Drosophila* species, which are adapted to oviposit in fermenting fruit of no value, *D. suzukii* has evolved a distinct ecological niche ^13,14^. It targets intact, ripening fruit for oviposition, causing significant agricultural losses^15^. This shift allows the species to exploit a previously underutilized resource, but it also poses new sensory and behavioral challenges. Navigating this niche requires the ability to detect and respond to a different set of volatile cues than those used by species that specialize on rotting substrates. Consequently, *D. suzukii*’s unique host preference is likely underpinned by extensive modifications to its olfactory system. Several molecular mechanisms are known to drive such adaptations, including amino acid substitutions that alter receptor tuning properties, changes in gene expression levels that modify receptor neuron sensitivity, and structural changes in the genome that increase or decrease receptor copy number through duplication, neofunctionalization, or pseudogenization ^5,6,16–20^.

Comparative genomic and transcriptomic analyses have revealed that *D. suzukii*’s odorant receptor (Or) repertoire exhibits distinctive evolutionary signatures ^21–24^. Two gene expansions, *Or23a* and *Or67a*, set this species apart from its close relatives *D. biarmipes* and *D. melanogaster*. In *D. suzukii*, *Or23a* is present in four intact copies (*Or23a1*, *Or23a2*, *Or23a3*, and *Or23a4*), while *Or67a* is represented by four functional paralogs (*Or67a1*, *Or67a2*, *Or67a3*, and *Or67a4*) and one pseudogene (*Or67a5*). Interestingly, only the *Or67a* expansion is retained in *D. biarmipes* ^22^, suggesting that the expansion of *Or23a* occurred more recently in the *D. suzukii* lineage. In *D. melanogaster*, *Or23a* is expressed in the B neuron of antennal intermediate 2 (ai2) sensilla, but extensive functional screening has not identified strong ligands, leaving its role unresolved ^25^.

*Or67a*, by contrast, is better functionally characterized. In *D. melanogaster*, *Or67a* is expressed in the A neuron of ab10 sensilla, where it mediates strong responses to aromatic esters such as benzyl butyrate, benzyl acetate, and phenethyl propionate ^25–27^. In *D. suzukii*, two copies of *Or67a* (*Or67a3* and *Or67a4*) exhibit signatures of positive selection ^22^, suggesting adaptive evolution potentially linked to the detection of odorants associated with ripening fruit.

Concurrently, *D. suzukii* has lost several ancestral Or genes, including *Or74a*, *Or85a*, and *Or98b* ^22,23^. These losses could represent either relaxed selection for receptors tuned to volatiles no longer relevant in the new ecological context, or functional trade-offs associated with receptor repertoire remodeling.

Despite these striking genomic changes, the extent to which receptor gains and losses have reshaped the peripheral olfactory architecture, altering neuron identities, response properties, and overall coding capacity, remains largely unexplored.

Here, through a combination of electrophysiological recordings and comparative analyses, we identified olfactory receptor neurons whose response profiles differ markedly between *Drosophila suzukii* and its close relatives, *D. melanogaster* and *D. biarmipes*. Using molecular genetics, CRISPR/Cas9 gene editing, AlphaFold, and molecular docking, we elucidated the molecular mechanisms underlying these neuronal differences. Moreover, our work revealed that both gene losses and gains have shaped the olfactory repertoire of *D. suzukii*, driving its sensory adaptation and facilitating its ecological niche transition from fermenting to ripening fruits. These findings highlight how molecular changes at the receptor level translate into behavioral shifts that underpin the evolutionary success and pest status of *D. suzukii*.

## RESULTS

### Neurons whose responses differ between *D. suzukii* and *D. melanogaster*

To identify neurons that differ in their responses between *D. suzukii* and *D. melanogaster*, we compared odorant-evoked responses of olfactory receptor neurons (ORNs) in the antennal basiconic, intermediate, and trichoid sensilla of both species using single sensillum recordings (SSR). Modifications in the tuning of these neurons could result from gene births and deaths accompanying *D. suzukii*’s transition to ripe fruit^28^ and may underlie its novel specialization. This analysis complements our prior work on maxillary palp basiconic and antennal coeloconic sensilla in *D. suzukii* ^29,30^ and together provides a comprehensive functional atlas of ORN responses across all sensillum types.

For this analysis, we used a panel of 113 ecologically relevant odorants, each tested at a 10^−2^ dilution, including acids, esters, alcohols, aldehydes, ketones, lactones, and terpenes (Figure S1). This analysis generated a dataset of 3,503 odorant–ORN combinations in each species (Figure S1; Extended Data Files S1 and S2) (n = 3–6 per combination in *D. suzukii* and n = 3 per combination in *D. melanogaster*) and revealed that the odorant response profiles of most ORNs are qualitatively similar between the two species(Figure S1). However, a subset of four ORNs displayed notable shifts in odorant tuning (Figure 1A). These ORNs are ab2B, ab3A, ab4B, and ab10A.

**Figure 1.**
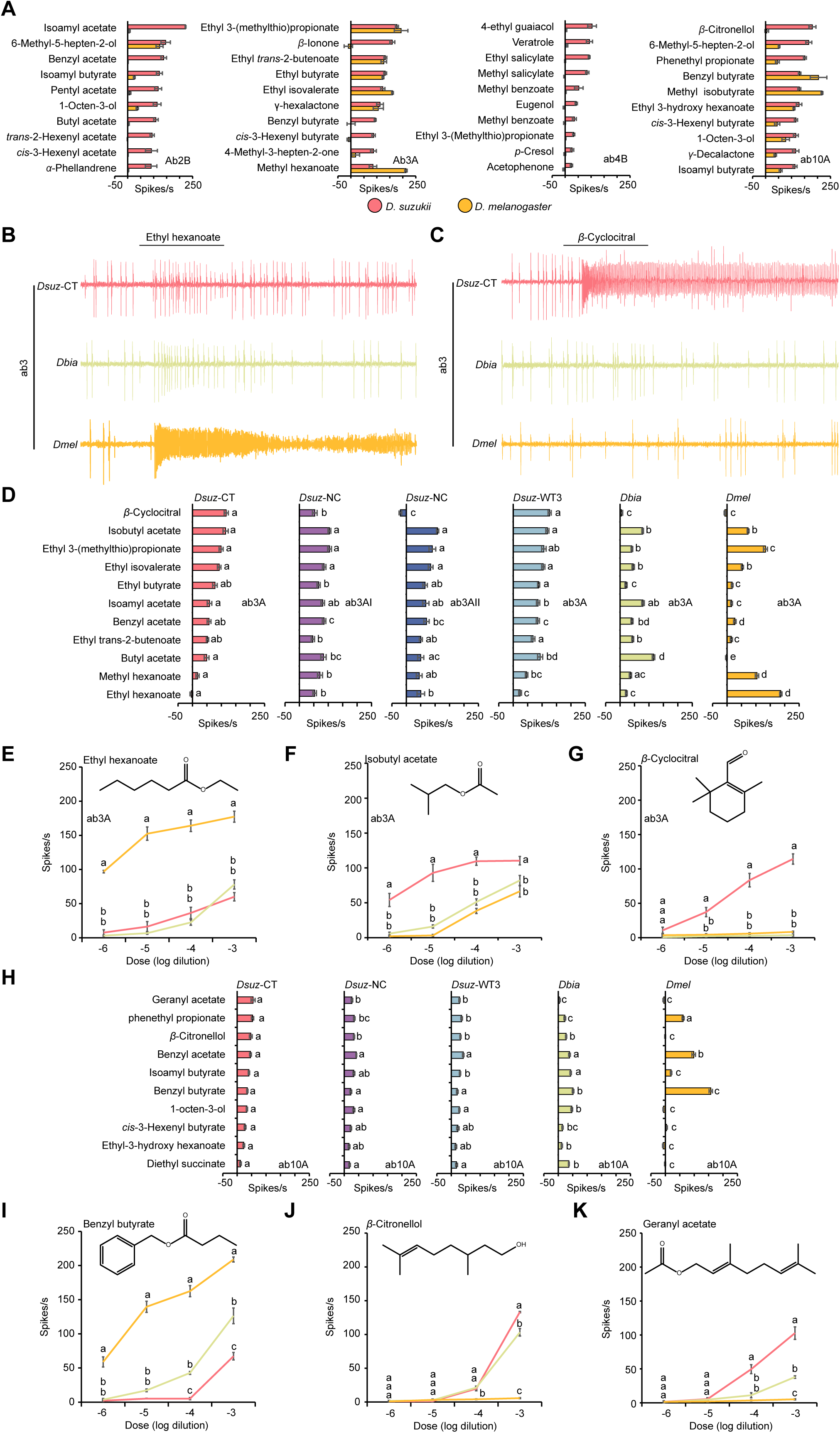
Shared shifts in odor tuning of ab3A and ab10A in *D. suzukii* and *D. biarmipes*. ((A) Responses of ab2B, ab3A, ab4B, and ab10A neurons in *D. suzukii* (n = 3–6) and *D. melanogaster* (n = 3) to the indicated odorants, each tested at a 10⁻² dilution. Error bars represent SEM. Responses to additional odorants are shown in Figure S1 and Extended Data Files S1 and S2. (B) Example traces of the response of the ab3 sensillum type to a 10^-4^ dilution of ethyl hexanoate in *D. suzukii* (CT-strain), *D. biarmipes*, and *D. melanogaster*. (C) Example traces of the response of the ab3 sensillum type to a 10^-4^ dilution of β-cyclocitral in *D. suzukii* (CT-strain), *D. biarmipes*, and *D. melanogaster*. (D) Responses of ab3A neurons in three strains of *D. suzukii* (*Dsuz-CT*, *Dsuz-NC*, and *Dsuz*-WT3), *D. biarmipes*, and *D. melanogaster* to a 10^-4^ dilution of each of the indicated odorants. One-way ANOVA followed by Tukey’s multiple comparison test; n = 10. Values marked with different letters are significantly different. Error bars represent SEM. (E-G) Responses of ab3A neurons to different dilutions of ethyl hexanoate (E), isobutyl acetate (F), and β-cyclocitral (G) in *D. suzukii* (CT-strain), *D. biarmipes*, and *D. melanogaster*. One-way ANOVA followed by Tukey’s multiple comparison test; n = 5 for each dilution. Values indicated with different letters are significantly different. Error bars represent SEM. (H) Responses of ab10A neurons in three strains of *D. suzukii*, *D. biarmipes*, and *D. melanogaster* to a 10^-4^ dilution of each of the indicated odorants. One-way ANOVA followed by Tukey’s multiple comparison test; n = 10. Values indicated with different letters are significantly different. Error bars represent SEM. (I-K) Responses of ab10A neurons to different dilutions of benzyl butyrate (I), β-citronellol (J), and geranyl acetate (K) in *D. suzukii* (CT-strain), *D. biarmipes*, and *D. melanogaster*. One-way ANOVA followed by Tukey’s multiple comparison test; n = 5 for each dilution. Values indicated with different letters are significantly different. Error bars represent SEM.

To validate and further dissect these interspecific differences, we performed a second round of SSR using an odorant panel that included the most potent ligands identified in our initial screen, together with ligands previously reported for the corresponding ORNs^31,32^. All odorants were tested at a 10^−4^ dilution to approximate naturalistic concentrations encountered by flies in the wild^33^.

To differentiate between *D. suzukii*-specific changes and more broadly shared evolutionary trends, we included *D. biarmipes* in our analysis. This species is the closest extant relative of *D. suzukii* ^34,35^ and has no distinct preference for either overripe or ripe fruit ^13,14,21^.

To assess the robustness of these shifts within *D. suzukii*, we included two geographically distinct populations: the WT3 strain from California (*Dsuz*-WT3), which served as the reference genome strain ^34^, and an independent strain collected in North Carolina (*Dsuz*-NC).

### Shared and species-specific shifts in *D. suzukii*

In ab3A neurons, we observed two kinds of changes, one of which is shared by *D. suzukii* and *D. biarmipes*. These included markedly reduced responses to volatiles that strongly activate this neuronal type in *D. melanogaster* (Figure 1B, D, E), including ethyl hexanoate, which is a fermentation product^31^, and increased responses to an alternative set of esters, including isoamyl acetate and butyl acetate (Figure 1D), which are produced during fruit ripening^31^.

The second kind of change is unique to *D. suzukii*. These included significantly stronger responses to isobutyl acetate, ethyl butyrate, and ethyl isovalerate compared to *D. biarmipes* and *D. melanogaster* (Figure 1D, F), as well as robust, dose-dependent responses to β-cyclocitral (Figure 1C, D, G), a volatile isoprenoid emitted by green foliage, including strawberry leaves^31^. By contrast, over a range of dilutions, this compound elicited little or no activity in the ab3A neurons of *D. biarmipes* and *D. melanogaster* (Figure 1C, D, G).

We also found intraspecific variation in responses to β-cyclocitral across *D. suzukii* strains. All ab3A neurons in the WT3 strain (California origin) responded robustly to β-cyclocitral, whereas responses in the North Carolina strain were more heterogeneous: some ab3A neurons responded strongly while others showed little or no response (Figure 1D).

In ab10A neurons, both *D. suzukii* and *D. biarmipes* exhibited reduced sensitivity to benzyl butyrate and benzyl acetate compared to *D. melanogaster* (Figure 1H, I) and enhanced responses to geranyl acetate, β-citronellol, isoamyl butyrate, 1-octen-3-ol, ethyl-3-hydroxyhexanoate, and diethyl succinate (Figure 1H, J, K).

In ab2B neurons, both *D. melanogaster* and *D. biarmipes* exhibited strong to ethyl 3-hydroxybutyrate, a compound closely associated with fermentation^36,37^, whereas all three *D. suzukii* strains displayed strongly reduced sensitivity (Figure 2A, C, D). By contrast, ab2B neurons in *D. suzukii* responded robustly to a distinct set of esters, including isoamyl acetate, butyl acetate, pentyl acetate, benzyl acetate, and isoamyl butyrate (Figure 2B, C, E, F). These esters are formed as part of the fruit ripening process^31,36,37^.

**Figure 2.**
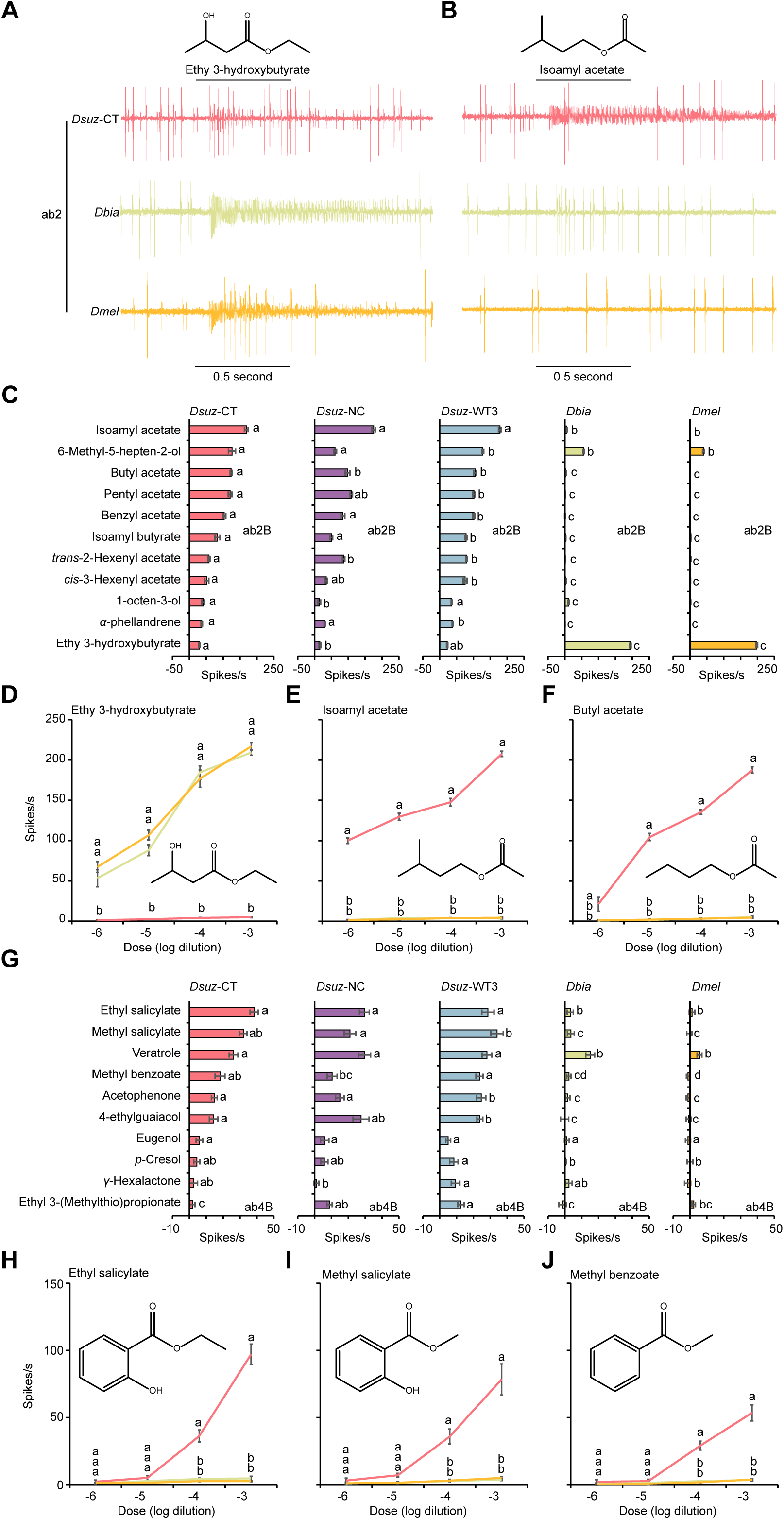
Species-specific shifts in odor tuning of ab2B and ab4B in *D. suzukii*. (A) Example traces of the response of the ab2 sensillum type to a 10^-4^ dilution of ethyl 3-hydroxybutyrate in *D. suzukii* (CT-strain), *D. biarmipes*, and *D. melanogaster*. (B) Example traces of the response of the ab2 sensillum type to a 10^-4^ dilution of isoamyl acetate in *D. suzukii* (CT-strain), *D. biarmipes*, and *D. melanogaster*. (C) Responses of ab2B neurons in three strains of *D. suzukii*, *D. biarmipes*, and *D. melanogaster* to a 10^-4^ dilution of each of the indicated odorants. One-way ANOVA followed by Tukey’s multiple comparison test; n = 10. Values marked with different letters are significantly different. Error bars represent SEM. (D-F) Responses of ab2B neurons to different dilutions of ethyl 3-hydroxybutyrate (D), isoamyl acetate (E), and butyl acetate (F) in *D. suzukii* (CT-strain), *D. biarmipes*, and *D. melanogaster*. One-way ANOVA followed by Tukey’s multiple comparison test; n = 5 for each dilution. Values indicated with different letters are significantly different. Error bars represent SEM. (G) Responses of ab4B neurons in three strains of *D. suzukii*, *D. biarmipes*, and *D. melanogaster* to a10^-4^ dilution of the indicated odorants. One-way ANOVA followed by Tukey’s multiple comparison test; n = 10. Values indicated with different letters are significantly different. Error bars represent SEM. (H-J) Responses of ab4B neurons to different dilutions of ethyl salicylate (H), methyl salicylate (I), and methyl benzoate (J) in *D. suzukii* (CT-strain), *D. biarmipes*, and *D. melanogaster*. One-way ANOVA followed by Tukey’s multiple comparison test; n = 5 for each dilution. Values indicated with different letters are significantly different. Error bars represent SEM.

In ab4B neurons, all three species responded to veratrole, whereas all three *D. suzukii* strains additionally responded to ethyl salicylate, methyl salicylate, methyl benzoate, acetophenone, and 4-ethylguaiacol (Figure 2G–J).

Together, these findings reveal a mosaic of shared and species-specific olfactory changes in *D. suzukii*. The shifts in ab3A and ab10A, shared with *D. biarmipes*, likely reflect phylogenetic divergence, whereas the unique sensitivity of ab3A neurons to β-cyclocitral and species-specific tuning in ab2B and ab4B neurons point to derived adaptations supporting *D. suzukii*’s specialization on ripe fruit.

### *D. suzukii* ab3A neurons co-express two odorant receptors, one of which is specifically tuned to β-cyclocitral

We next focused on ab3A neurons, which display a robust response to the strawberry leaf volatile β-cyclocitral. We hypothesized that this neuronal type in *D. suzukii* co-expresses two receptors, one of which is specifically tuned to β-cyclocitral. This hypothesis was based on three observations: (1) only *D. suzukii* ab3A neurons respond robustly to β-cyclocitral, despite both *D. suzukii* and *D. biarmipes* having a single functional gene (referred to as the *Or22a*-like receptor) at the locus that determines ab3A responses in drosophilids, with 91% sequence identity between the two species; (2) the North Carolina strain of *D. suzukii* harbors two ab3A populations, one responsive to β-cyclocitral and one not; and (3) the chemical structure of β-cyclocitral, a cyclic isoprenoid aldehyde, differs markedly from the long- and short-chain esters that typically activate ab3A neurons in *D. suzukii* and other *Drosophila* species.

To test this hypothesis, we first expressed the *D. suzukii Or22a* (*DsuzOr22a*) in the *D. melanogaster* ab3A “empty neuron” system ^38^, where the endogenous *Or22a*/*Or22b* locus is deleted and replaced by *Gal4* (Figure 3A). Ectopic expression of *DsuzOr22a* restored responses to a suite of esters that matched the tuning profile of native *D. suzukii* ab3A neurons (Figure 3B, D). However, these neurons did not confer responses to β-cyclocitral (Figure 3C, D), indicating that *DsuzOr22a* is not sufficient for β-cyclocitral detection and that a second co-expressed receptor is required.

**Figure 3.**
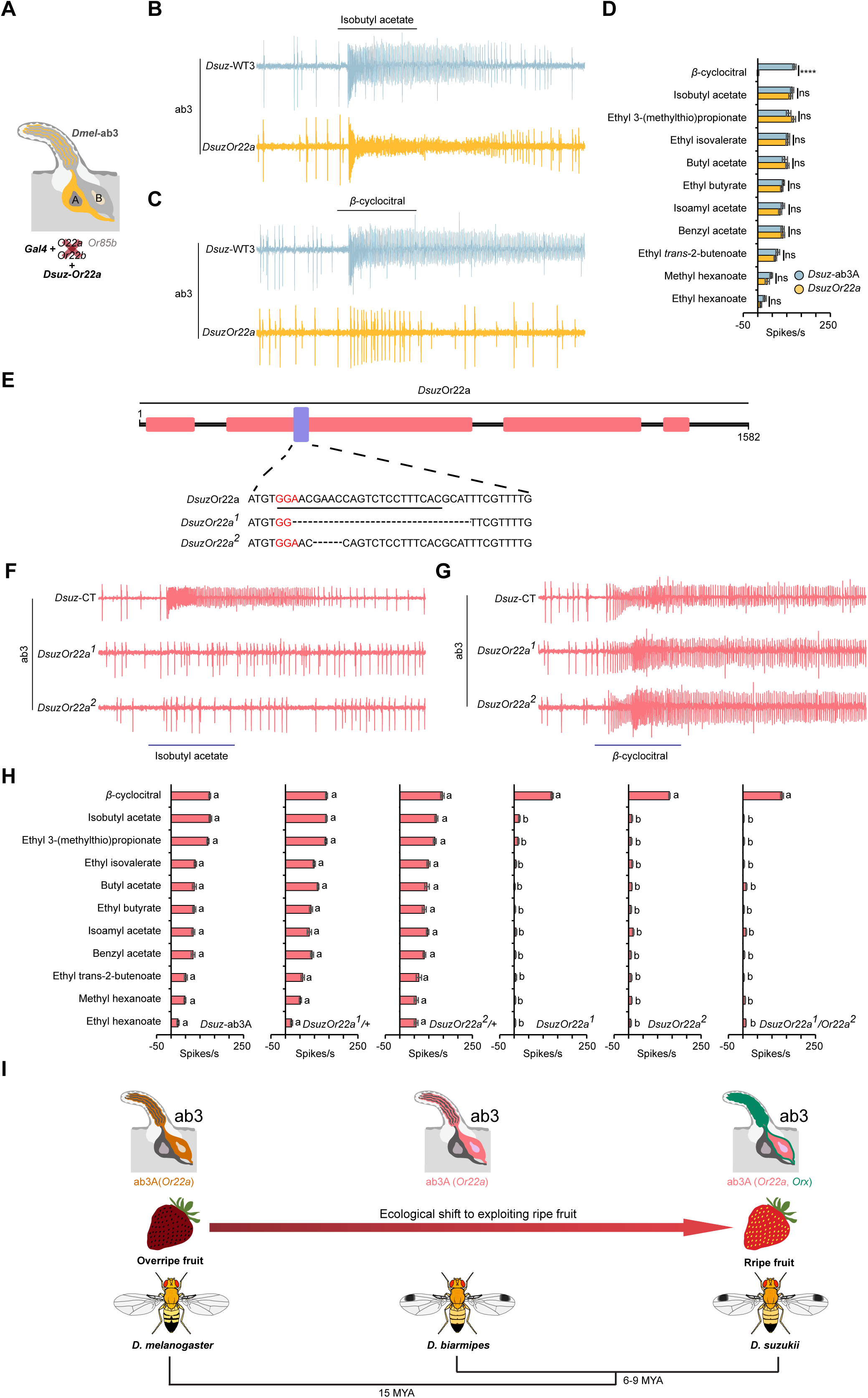
ab3A neurons in *D. suzukii* express two odorant receptors, one of which is tuned to β-cyclocitral. (A) Illustration of the *D. melanogaster* empty neuron system. (B) Example traces of the responses of *D. suzukii* ab3A neurons and *DsuzOr22a* to a 10^-4^ dilution of isobutyl acetate. (C) Example traces of the responses of *D. suzukii* ab3A neurons and *DsuzOr22a* to a 10^-4^ dilution of β-cyclocitral. (D) Responses of *D. suzukii* ab3A neurons and *DsuzOr22a* to eleven odorants, each tested at a 10^-4^ dilution. ns = not significant; ****p<0.0001; Mann-Whitney test; n = 10. Error bars represent SEM. (E) Schematic illustrating the generation of *DsuzOr22a* mutants using CRISPR/Cas9. Nucleotides in pink represent the PAM site, underlined nucleotides indicate the gRNA sequence, and dashed lines denote deleted nucleotides. (F) Example traces of the responses of ab3A neurons in wild-type *D. suzukii*, *DsuzOr22a^1^,* and *DsuzOr22a^2^* to a 10^-4^ dilution of isobutyl acetate. (G) Example traces of the responses of ab3A neurons in wild-type *D. suzukii*, *DsuzOr22a^1^,* and *DsuzOr22a^2^* to a 10^-4^ dilution of β-cyclocitral. (H) Responses of ab3A neurons in wild-type *D. suzukii*, *DsuzOr22a^1/+^, DsuzOr22a^2/+^, DsuzOr22a^1^,* and *DsuzOr22a^2^* to a panel of eleven odorants, each tested at a 10^-4^ dilution. One-way ANOVA followed by Tukey’s multiple comparison test; n = 10. Values indicated with different letters are significantly different. Error bars represent SEM. (I) Diagrams of ab3 sensilla in *D. melanogaster*, *D. biarmipes*, and *D. suzukii*. In *D. melanogaster* and *D. biarmipes*, the ab3A neuronal type expresses a single odorant receptor (*Or22a*), whereas in *D. suzukii* it co-expresses two receptors (*Or22a* and *Orx*).

We next used CRISPR/Cas9-mediated mutagenesis to generate loss-of-function alleles of *DsuzOr22a*. Two independent alleles were recovered: *DsuzOr22a^1^*, carrying a 25-nt deletion, and *DsuzOr22a^2^*, carrying a 4-nt deletion. Both mutations are predicted to produce truncated, non-functional proteins of 106 amino acids (Figure 3E). To control for potential background effects, we also generated heteroallelic *DsuzOr22a* mutant combinations using these homozygous lines.

Compared to Wild-type, *DsuzOr22a^1/+^*, and *DsuzOr22a^2/+^* flies, ab3A neurons in *DsuzOr22a^1^, DsuzOr22a^2^, and DsuzOr22a^1/^Or22a^2^* mutant flies exhibited drastically reduced responses to all tested esters (Figure 3F, H), demonstrating that *DsuzOr22a* is required for ester sensitivity. However, responses to β-cyclocitral were intact in all mutants, confirming that a second receptor mediates detection of this compound (Figure 3G, H).

To determine whether the β-cyclocitral receptor belongs to the odorant receptor (OR) or ionotropic receptor (IR) family, we measured responses from large basiconic sensilla in *D. suzukii Orco* mutant flies. Because *Orco* is an obligate co-receptor for all functional ORs but not IRs, the presence or absence of β-cyclocitral responses in this background would distinguish the receptor class.

We found that β-cyclocitral did not elicit any response in any of the 117 individual sensilla examined in *DsuzOrco* mutant flies, indicating that β-cyclocitral detection depends on *Orco* and thus is mediated by an OR rather than an IR (Figure S2).

Together, these results demonstrate that *D. suzukii* ab3A neurons co-express two odorant receptors (Figure 3I): *Or22a*, which has shifted its tuning toward several ripening fruit esters such as isobutyl acetate, butyl acetate, and isoamyl acetate, and a second, as-yet-unidentified receptor that is specifically tuned to β-cyclocitral.

### A Single Amino Acid Substitution Alters Ligand Specificity of *Or22a* in *D. suzukii*

We next investigated the molecular basis of the tuning shift of *DsuzOr22a*. To do this, we aligned the *Or22a* amino acid sequences from *D. melanogaster*, *D. suzukii*, and *D. biarmipes* using Clustal Omega and visualized the alignment with MView (Figure 4A). This analysis revealed 97 amino acid substitutions shared between *D. suzukii* and *D. biarmipes* relative to *D. melanogaster*, including a substitution at position 201 (aspartic acid [D] to alanine [A]). In *D. melanogaster* strains, the residue at this position is predicted to lie within the odorant-binding pocket^39^, which is proposed to form within the extracellular leaflet of the plasma membrane between helices TM1–TM6^40^. Previous work showed variation at this site, together with changes at residue 92, alters ligand-binding specificity and contributes to phenotypic variation in ab3A responses among natural populations of *D. melanogaster*^40^. Since residue 92 is conserved across all three species, the substitution at position 201 likely plays a major role in the tuning shift of *DsuzOr22a*.

**Figure 4.**
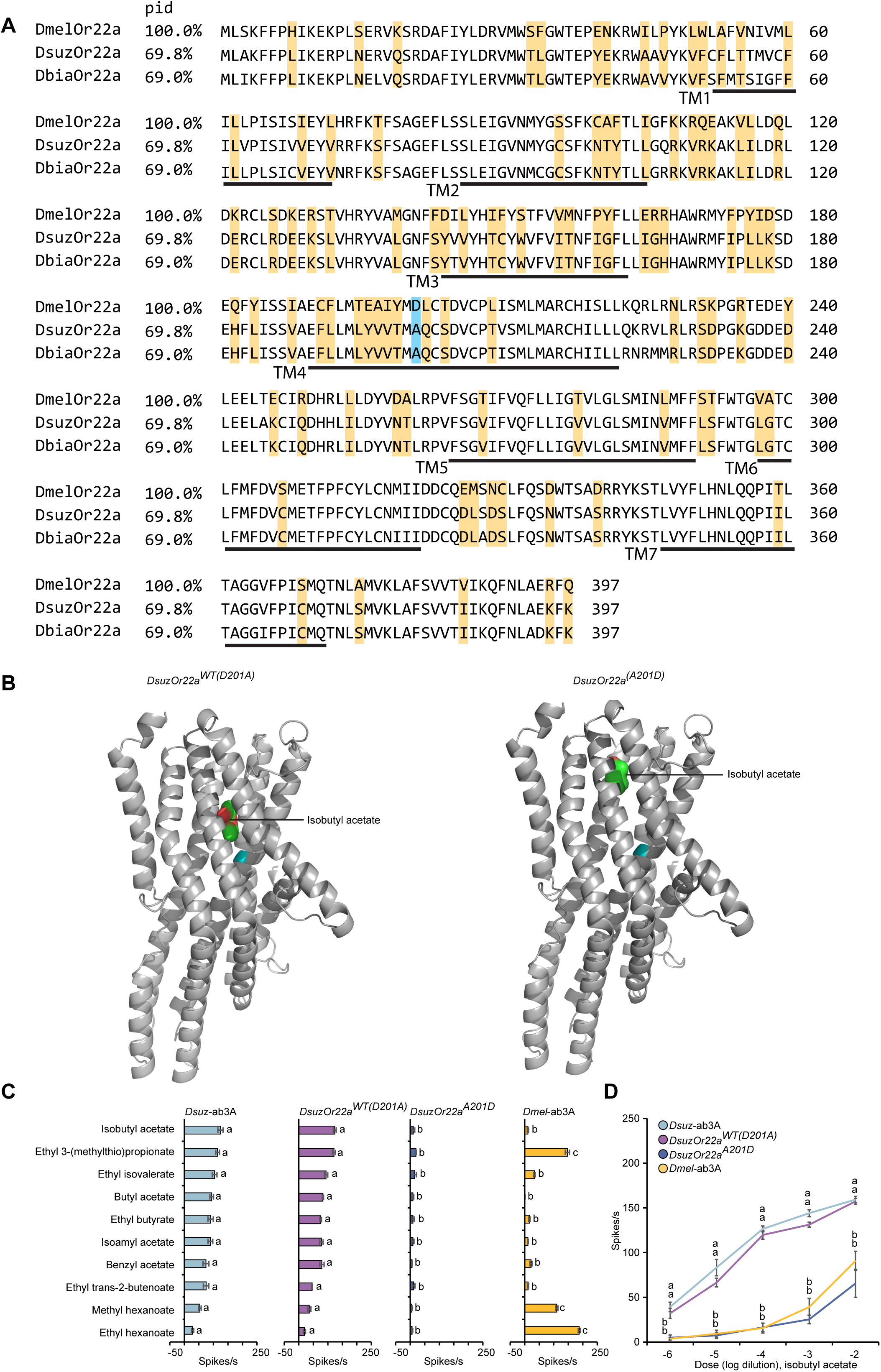
A single amino acid substitution alters the architecture of the ligand-binding pocket of Or22a in *D. suzukii*. (A) Protein sequence alignment of Or22a orthologs of *D. melanogaster*, *D. suzukii*, and *D. biarmipes*. Predicted transmembrane (TM) domains are indicated with black lines. Yellow shading highlights amino acids conserved between *D. suzukii* and *D. biarmipes* but differing from D. melanogaster. Cyan shading marks the amino acid at position 201 in all three species. PID indicates percent sequence identity. (B) Cartoon models of *DsuzOr22a^WT(D201A)^*(left) and *DsuzOr22a^(A201D)^* (right) in lateral view, showing the predicted binding sites with docked isobutyl acetate. Position 201 is highlighted in cyan. (C) Responses of wild-type *D. suzukii* ab3A neurons, *DsuzOr22a^WT(D201A)^, DsuzOr22a^(A201)^*, and wild-type *D. melanogaster* ab3A neurons to a panel of ten odorants, each tested at a 10^-4^ dilution. One-way ANOVA followed by Tukey’s multiple comparison test; n = 10. Values indicated with different letters are significantly different. Error bars represent SEM. (D) Responses of wild-type *D. suzukii* ab3A neurons, *DsuzOr22a^WT(D201A)^, DsuzOr22a^(A201)^*, and wild-type *D. melanogaster* ab3A neurons to different dilutions of isobutyl acetate. One-way ANOVA followed by Tukey’s multiple comparison test; n = 10. Values indicated with different letters are significantly different. Error bars represent SEM.

To determine the functional impact of this substitution, we used AlphaFold2 to model the wild-type DsuzOr22a (D201A) and an *in silico* variant with the A201D substitution. We then performed molecular docking with isobutyl acetate, a key ligand for *DsuzOr22a*. The predicted models revealed that the A201D substitution induces a pronounced reorganization of the binding pocket architecture, producing a conformation distinct from that of the wild-type receptor (Figure 4B). This suggests that D201A significantly alters receptor–ligand interactions and contribute to the tuning shift of *DsuzOr22a*.

We then tested this prediction by generating a site-directed mutant of *DsuzOr22a* (A201D) and expressing it in the *D. melanogaster* ab3A “empty neuron” system. This substitution did not increase the response of *DsuzOr22a* to any of the compounds that strongly activate *D. melanogaster* ab3A (Figure 4C). By contrast, it reduced responses to all tested odorants (Figure 4C, D). Interestingly, responses of the A201D variant to isobutyl acetate across a range of dilutions more closely resembled that of *D. melanogaster* ab3A than of *D. suzukii* ab3A or wild-type *DsuzOr22a* (Figure 4D), indicating a partial reversion toward the ancestral tuning state. These results support a critical role for residue 201 in reshaping *Or22a* ligand specificity in *D. suzukii*, particularly for fruit-ripening esters.

### Functional Analysis of *Or23a* and *Or67a* Paralogs in *D. suzukii*

Having demonstrated that the response to β-cyclocitral in *D. suzukii* ab3A depends on an odorant receptor other than *Or22a* and deciphered the molecular basis of the *DszuOr22a* tuning shift, we next investigated the contributions of the expanded *Or23a* (four paralogs) and *Or67a* (five paralogs) receptor subfamilies to β-cyclocitral sensitivity in ab3A neurons, as well as to tuning shifts in ab10A and ab2B neurons. To do this, we expressed each paralog individually in the *D. melanogaster* ab3A “empty neuron” system and measured odor-evoked responses across a broad panel of compounds.

We found that all *Or23a* paralogs showed little (<30 spikes/s) or no response to any of the tested odorants (Extended Data File S3), indicating that their key ligands remain unidentified. Similarly, *Or67a5* did not respond to any odorant, consistent with its annotation as a pseudogene due to a premature stop codon (TAA) at nucleotide position 174 (Extended Data File S4).

By contrast, *Or67a1*, *Or67a2*, *Or67a3*, and *Or67a4* each exhibited robust and specific responses to multiple odorants (Figures 5). However, none of these receptors responded to β-cyclocitral with the same magnitude or sensitivity as the native ab3A neuron, indicating that the receptor mediating this response remains unidentified.

**Figure 5.**
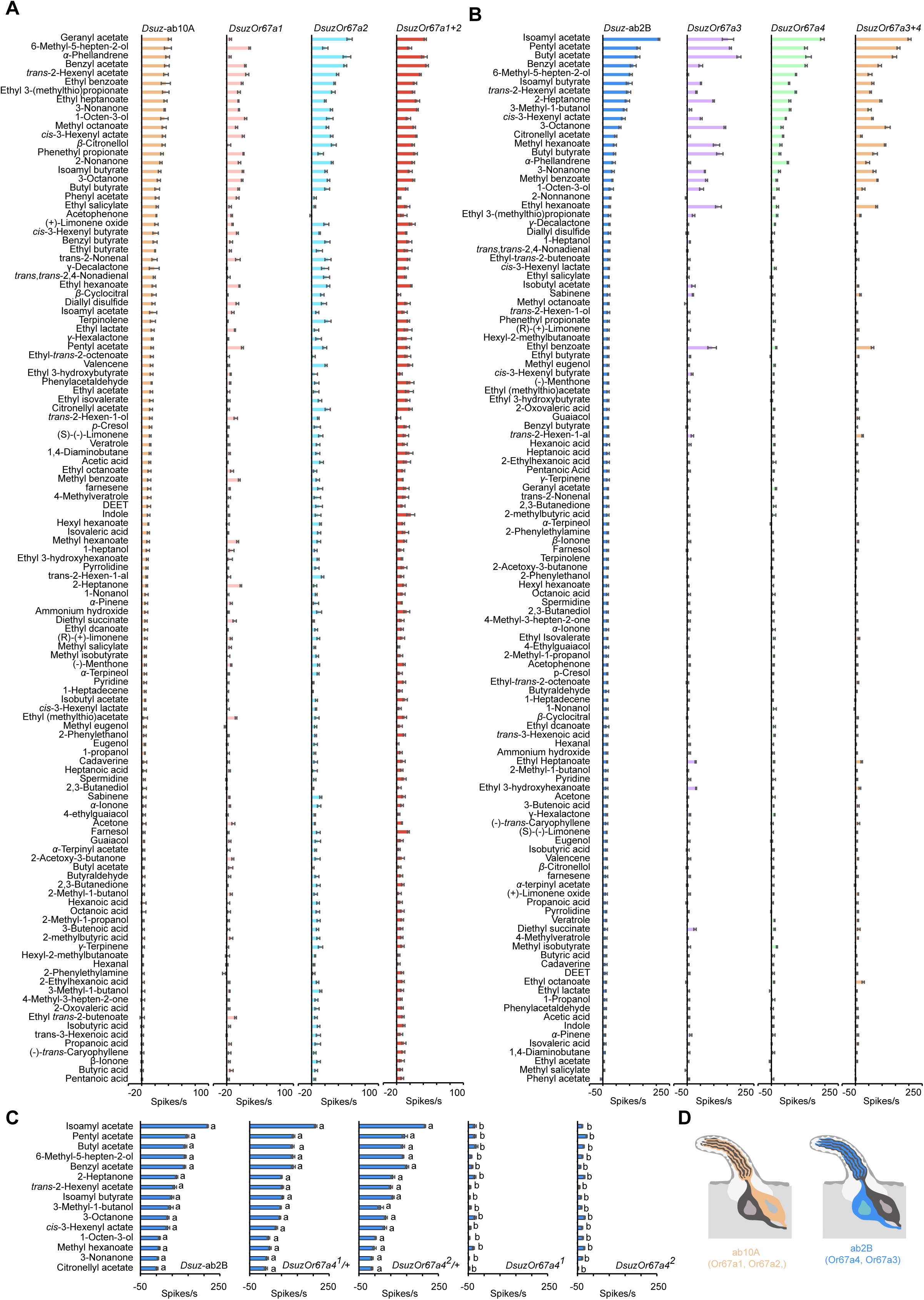
Responses of *D. suzukii Or67a* paralogs to a battery of 119 odorants. (A) Responses of *Dsuz*-ab10A, *DsuzOr67a1*, *DsuzOr67a2*, and *DsuzOr67a1+2* to a battery of 119 odorants, each tested at a 10-4 dilution. n = 5. Error bars represent SEM. (B) Responses of *Dsuz*-ab2B, *DsuzOr67a3*, *DsuzOr67a4*, and *DsuzOr67a3+4* to a battery of 119 odorants, each tested at a 10-4 dilution. n = 5. Error bars represent SEM. (C) Responses of ab2B neurons in wildtype *D. suzukii*, *DsuzOr67a^1/+^, DsuzOr67a^2/+^, DsuzOr67a^1^,* and *DsuzOr67a^2^* to a panel of 15 odorants, each tested at a 10^-4^ dilution. One-way ANOVA followed by Tukey’s multiple comparison test; n = 10. Values indicated with different letters are significantly different. Error bars represent SEM. (D) Diagrams of ab2 and ab10 sensilla showing the expression of Or67a paralogs in ab10A and ab2B neurons of *D. suzukii*.

### Co-expression of *Or67a1* and *Or67a2* Produces a Response Profile Closely Matching the Native ab10A Odor Response in *D. suzukii*

*Or67a1* and *Or67a2* responded to an overlapping set of odorants, but differed in response magnitude for several compounds, including geranyl acetate, α-phellandrene, and β-citronellol (Figure 5A). The odorant response profile of *Or67a2* more closely resembled that of the native *D. suzukii* ab10A neurons than did *Or67a1* (Figure 5A), with a mean Euclidean distance of 116 ± 0.8 (SE) compared to 122 ± 1 for *Or67a1*.

To determine whether co-expression of both receptors is required to fully replicate the ab10A response profile, we expressed *Or67a1* and *Or67a2* together in the *D. melanogaster* ab3A “empty neuron” system. This co-expression produced responses that closely matched the ab10A response profile (Figure 5A), with a smaller Euclidean distance (107 ± 2.5) than either receptor alone. The simple interpretation of these results is that *Or67a1* and *Or67a2* are co-expressed in the *D. suzukii* ab10A neurons and that their combined activity produces the characteristic tuning of this neuronal type.

### *Or67a4* is expressed in the ab2B neurons of *D. suzukii* and mediates their odorant responses

*Or67a3* and *Or67a4*, on the other hand, displayed odorant response profiles that diverged markedly from that of ab10A neurons (Figure 5B). Surprisingly, the response profile of each of *Or67a3* and *Or67a4* closely resembled that of the native *D. suzukii* ab2B neurons (Figure 5B), with Or67a4 showing a smaller mean Euclidean distance (180 ± 2.5) compared to Or67a3 (319 ± 2.7). However, co-expression of *Or67a3* and *Or67a4* in the *D. melanogaster* ab3A ‘empty neuron’ system produced a response profile more similar to *Or67a3* (192 ± 1.4 SE, mean Euclidean distance) than to either the native *D. suzukii* ab2B (229 ± 2.5) or *Or67a4* alone (218 ± 1.1) (Figure 5B). The simple interpretation of these results is that both *Or67a3* and *Or67a4* are expressed in ab2B neurons, since no other ORNs besides ab2B produce responses resembling those of *Or67a3*, with *Or67a4* dominating the overall odorant response profile in the native ab2B, likely due to higher expression levels. Consistent with this, *Or67a4* expression increases significantly after mating, while Or67a3 expression remains unchanged^41^.

We further confirmed and extended these findings by generating two independent loss-of-function alleles of *DsuzOr67a4* using CRISPR/Cas9: one with two deletions (22 bp and 47 bp; *DsuzOr67a4^1^*) (*DsuzOr67a4^2^*) and the other with a 22-bp deletion (Figure S3). From these homozygous mutants, we also established two heterozygous lines.

While wild-type and heterozygous lines responded indistinguishably, ab2B neurons in the two *DsuzOr67a4* mutant alleles exhibited markedly reduced responses to all tested odorants (Figure 5C). This indicates that *Or67a4* is expressed in *D. suzukii* ab2B neurons and is essential for their odorant responses. Alternatively, because *Or67a4* lies only 380 bp upstream of *Or67a3* and the two genes appear to undergo transcriptional readthrough (Figure S3C), its deletion could disrupt shared regulatory elements or chromatin structure, thereby altering *Or67a3* transcription. The phenotype may thus reflect both *Or67a4* loss and secondary effects on O*r67a3*.

Together, these findings reveal a clear functional and spatial partitioning among the *Or67a* paralogs in *D. suzukii*. *Or67a1* and *Or67a2* are co-expressed in ab10A neurons, while *Or67a4* and *Or67a3* are expressed in a separate neuronal type, ab2B. This partitioning of expression suggests that gene duplication in the *Or67a* lineage has facilitated subfunctionalization across distinct olfactory neuron types, enabling *D. suzukii* to broaden its olfactory repertoire and fine-tune its detection of ecologically relevant odorants.

### Species-Specific Partitioned Expression of Or67a Paralogs in Drosophila suzukii

To determine whether the evolutionary expansion of the *Or67a* subfamily in *D. suzukii* is associated with species-specific adaptations, we first inferred a maximum likelihood protein tree for the five *D. suzukii* paralogs (*Or67a1*, *Or67a2*, *Or67a3*, *Or67a4*, and *Or67a5*), the four *D. biarmipes* paralogs (*Or67a1*, *Or67a2*, *Or67a3*, and *Or67a5*), and *D. melanogaster Or67a*. We confirmed the absence of *Or67a4* in *D. biarmipes* through genome inspection and BLAST searches of the NCBI database using the *D. suzukii Or67a4* sequence.

This analysis revealed that the *Or67a* subfamily in *D. suzukii* is shaped by both conservation and lineage-specific innovation. *D. suzukii* and *D. biarmipes Or67a5* clustered with *D. melanogaster Or67a* with strong support (96 bootstrap) (Figure 6A), identifying it as the conserved ancestral ortholog that became pseudogenized in *D. suzukii*. *D. suzukii Or67a3* and *D. biarmipes Or67a3* clustered with *D. suzukii Or67a4* with full support (100 bootstrap) (Figure 6A), indicating that *Or67a4* originated through a lineage-specific duplication and divergence event in *D. suzukii*. By contrast, *D. suzukii Or67a1* and *Or67a2* each clustered with their *D. biarmipes* orthologs (100 bootstrap) (Figure 6A).

**Figure 6.**
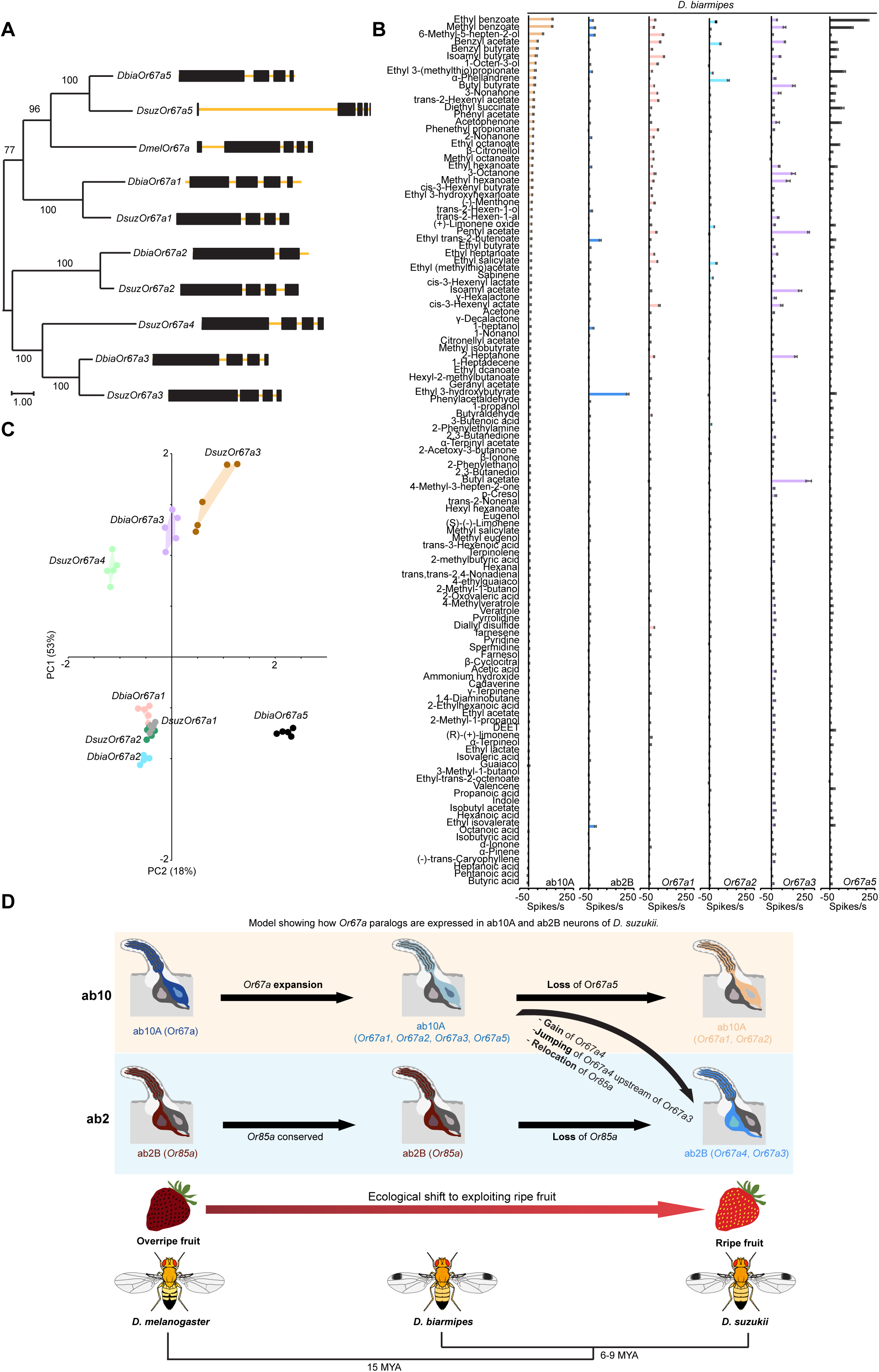
Evolution of the Or67a subfamily in *D. biarmipes*. (a) Maximum-likelihood protein tree inferred for *D. biarmipes* Or67a paralogs, *D. suzukii* Or67a paralogs, and *D. melanogaster* Or67a. Node values indicate bootstrap support based on 1,000 replicates. The scale bar represents the number of amino acid substitutions per site. Gene structures are depicted with black boxes for exons and yellow lines for introns. (b) Responses of *D. biarmipes* ab10A, ab2B, and *Or67a* paralogs to a battery of 119 odorants. n = 5. Error bars represent SEM. (C) Principal component analysis of the data from Figures 5A, 5B, and 6B. Percentages along the axes indicate the proportion of variance explained by each principal component. (D) Model showing how *Or67a* paralogs are expressed in ab10A and ab2B neurons of *D. suzukii*.

We next characterized the responses of *D. biarmipes Or67a* paralogs. Specifically, we asked whether the four intact *D. biarmipes* paralogs map onto the same neuronal classes (ab2B and ab10A) as in *D. suzukii* and whether their odor response profiles reflect conserved tuning or reveal species-specific divergence.

This analysis revealed that none of the *D. biarmipes Or67a* paralogs produced responses resembling those of ab2B neurons (Figure 6B; Extended Data File S5), indicating that the partitioned expression of *Or67a* paralogs between ab10A and ab2B is unique to *D. suzukii*.

*D. biarmipes Or67a1* and *Or67a2* responses closely resemble those of their *D. suzukii* orthologs (Figure 6B, C), reflecting both functional conservation and their close evolutionary relationships in the protein-based phylogenetic tree. However, *D. biarmipes Or67a2* responded to fewer compounds than its *D. suzukii* counterpart. These receptors appear to be expressed in ab10A neurons, as their responses closely match those of ab10A than of ab2B (Figure 6B).

*D. biarmipes Or67a5* showed strong responses to ethyl benzoate and methyl benzoate (Figure 6B), two of the best activators of the ancestral *Or67a* in *D. melanogaster*^27^. This functional conservation aligns with its clustering alongside *D. melanogaster Or67a* in the protein-based phylogenetic tree (Figure 6A). In *D. biarmipes*, ab10A neurons likely express this receptor, as these compounds are their strongest activators (Figure 6B). By contrast, the Or67a5 ortholog in *D. suzukii* is pseudogenized, and its ab10A neurons showed little or no response to these compounds (Figure 5A).

Most *Or67a3* responses are conserved between *D. biarmipes* and *D. suzukii* (Figure 6B, C), implying that this tuning profile evolved prior to the ecological shift of *D. suzukii* to exploit ripe fruit. *In D. suzukii*, *Or67a3* responses closely resemble those of ab2B (Figure 5B), whereas in *D. biarmipes*, *Or67a3* responses do not (Figure 6B), suggesting that this receptor likely remains expressed in ab10A neurons and that its functional contribution may be partially masked by co-expressed receptors within the same neuronal type. This is consistent with observations in taste neurons, where co-expression of multiple gustatory receptors can obscure individual contributions, and deletion of specific receptors often reveals novel responses^42^.

Altogether, these results support a model for the partitioning of Or67a paralog expression between two distinct neuronal types, ab10A and ab2B, in *D. suzukii* (Figure 6D). This scenario likely began with the ancestral expansion of *Or67a* into four paralogs (*Or67a1*, *Or67a2*, *Or67a3*, and *Or67a5*) in *D. biarmipes*, all of which were expressed in the ancestral ab10A neurons, with *Or67a1*, *Or67a2,* and *Or67a5* dominating the odorant response profile of this neuron type. In *D. suzukii*, the ecological shift toward exploiting ripe fruit coincided with the pseudogenization of *Or67a5* and the loss of *Or85a* expression in ab2B neurons. This loss was compensated by the emergence of *Or67a4*, which relocated to only 380 bp upstream of *Or67a3*. This genomic rearrangement likely altered regulatory elements, allowing both *Or67a3* and *Or67a4* to be expressed in ab2B neurons, thereby replacing *Or85a* and shaping the ab2B response profile of *D. suzukii* for fruit-ripening esters.

## DISCUSSION

Our results reveal that the evolutionary shift in preference from overripe to ripe fruit in *Drosophila suzukii,* a worldwide berry fly pest, is mirrored by functional changes in a limited subset of ORNs. Specifically, four ORNs (ab2B, ab3A, ab4B, and ab10A) exhibit altered odorant tuning in *D. suzukii* relative to *D. melanogaster*. Of these, ab3A and ab10A exhibit shared tuning shifts in *D. suzukii* and its non-pest sibling species *D. biarmipes*. These shared shifts likely reflect ancestral adaptations predating the transition to ripe fruit use. By contrast, ab2B and ab4B show unique tuning profiles in *D. suzukii*.

Importantly, these modifications are not solely the result of receptor sequence evolution but are also mediated by innovative mechanisms, including the co-expression of two distinct odorant receptors in a single ORN (as observed in ab3A) and the partitioned expression of gene duplicates across different ORNs (as seen with the *Or67a* paralogs in ab2B and ab10A). These strategies provide a flexible framework for expanding olfactory function without requiring the evolution of entirely new neuronal types, facilitating rapid and efficient sensory adaptation during ecological transitions.

### Receptor Co-expression Enables Novel Odor Tuning in ab3A of *D. suzukii*

The ab3A neurons of *D. suzukii* illustrate two distinct molecular innovations that align with its shift to ripe fruit. First, these neurons have lost responses to key fermentation volatiles that strongly activate ab3A in *D. melanogaster*. Instead, they have gained increased sensitivity to a new set of ripening-associated short-chain esters. Second, ab3A neurons in *D. suzukii* have acquired strong sensitivity to β-cyclocitral, a volatile isoprenoid commonly found in green plant tissues such as strawberry leaves^31^. This represents a major functional innovation, as neither *D. melanogaster* nor *D. biarmipes* ab3A neurons respond to this compound. Thus, detection of β-cyclocitral likely serves as a vegetative cue that signals the presence of a host plant bearing fruit, providing *D. suzukii* with a sensory advantage in locating oviposition sites while the fruit is still attached to the plant.

The capacity for functional plasticity in ab3A neurons is a recurring theme in *Drosophila* evolution. This neuronal type has repeatedly shifted its odorant tuning across species and even among populations, often in response to divergent ecological pressures^39,43–47^. These shifts are frequently mediated by amino acid changes in the *Or22* locus, which encodes the primary odorant receptor(s) in ab3A^20,45,47^. In the noni-specialist *D. sechellia*, for example, *Or22a* has acquired substitutions that enhance its sensitivity to methyl esters, which are abundant in noni fruit^45^. Genetic approaches have demonstrated that these sequence changes are sufficient to alter the ab3A sensitivity and that this receptor is important for mediating long-range attraction to noni^45^.

Similarly, in *D. suzukii*, we identified multiple amino acid substitutions in *Or22a*, including two at known functional hotspots, I45A and D201A, previously implicated in tuning modulation in other *Drosophila* species^20,45^. Our structural modeling and molecular docking analyses indicate that the D201A substitution plays a key role in the tuning shift of *Or22a* in *D. suzukii*. This substitution reorganizes the architecture of the ligand-binding pocket, likely altering the shape and electrostatic environment of the odorant interface. Furthermore, functional assays using a site-directed A201D mutant revealed markedly reduced responses to odorants, with its responses to isobutyl acetate across dilutions more closely resembling those of *D. melanogaster* ab3A than *D. suzukii* ab3A or wild-type *DsuzOr22a*, indicating partial reversion to the ancestral tuning state. However, similar tuning shifts and a substitution at position 201 were also observed in *D. biarmipes*, suggesting that *Or22a* diversification occurred prior to the shift to ripe fruit and may have evolved under different selective pressures.

Despite the functional impact of *Or22a* sequence evolution, it alone cannot account for the novel β-cyclocitral sensitivity. This response is absent in both *D. melanogaster* and *D. biarmipes*, indicating that the gain of β-cyclocitral sensitivity is a derived feature unique to *D. suzukii*. Our genetic and functional analyses revealed that β-cyclocitral sensitivity is mediated by a second odorant receptor co-expressed with *Or22a* in ab3A neurons. The response to β-cyclocitral was eliminated in *Orco* mutant flies confirming that the receptor that mediates this response belongs to the OR family rather than the ionotropic or gustatory receptor families. Recruitment of a second receptor likely reflects the chemical dissimilarity of β-cyclocitral from typical *Or22a* ligands and represents a strategic expansion of the odorant space encoded by a single ORN. This co-expression may also enable more refined odorant coding: combined input from both receptors indicates the presence of a high-quality oviposition site, while activation of *Or22a* alone signals suboptimal substrates. Such combinatorial coding could sharpen host selection and oviposition decisions.

### *Or67a* Gene Family Expansion and Deployment Across Neurons

The ab10A and ab2B neurons in *D. suzukii* have also undergone substantial functional and molecular remodeling. In *D. melanogaster*, ab10A neurons express a single receptor, *Or67a*, tuned to a broad range of compounds^25,26^, whereas ab2B neurons express *Or85a*, which responds to esters and alcohols typically associated with microbial fruit degradation^25–27,36^.

The response profiles of these two neuronal types have shifted markedly during *Drosophila* evolution^32,48,49^. Changes in ab10A are likely driven by *Or67a* gene expansion; for example, *D. simulans* carries three *Or67a* paralogs, all expressed in ab10A and contributing to its novel response profile^16^. By contrast, the mechanisms underlying ab2B tuning shifts remain less well understood. In several species, changes in ab2B odorant response profiles are associated with the loss of *Or85a*^48,49^, but the receptors that replace it remain unidentified.

In *D. suzukii*, *Or85a* is not detected in the genome^22,23^, while *Or67a* has expanded into five paralogs: four intact paralogs (*Or67a1* through *Or67a4*) and one pseudogene, *Or67a5*, which contains a premature stop codon^22,23^. Our functional characterization of these paralogs revealed a clear partitioning of activity. *Or67a1* and *Or67a2* are likely co-expressed in ab10A neurons, and their co-expression produced a response profile that is more similar to that of ab10A than either receptor alone. By contrast, Or67a4 and Or67a3 exhibited responses that are similar to those of ab2B neurons, and deletion of *Or67a4* severely reduced ab2B responses to odorants, providing direct evidence that this receptor underlies the novel tuning of ab2B neurons in *D. suzukii*.

*D. biarmipes* has also expanded *Or67a* into four functional paralogs (*Or67a1*, *Or67a2*, *Or67a3*, and *Or67a5*). Genome inspection and BLAST searches confirmed the absence of *Or67a4* in this species, indicating that this paralog is unique to *D. suzukii*. *In D. suzukii*, *Or67a4* relocated to 380 bp upstream of *Or67a3*, likely altering regulatory elements to enable co-expression of both receptors in ab2B neurons.

The tuning shift in *D. biarmipes* ab10A neurons is likely driven by co-expression of all *Or67a* paralogs, similar to *D. simulans*^16^. This conclusion is supported by several observations: First, none of the *D. biarmipes Or67a* paralogs produce responses resembling those of ab2B neurons, and ab2B neurons retain a *D. melanogaster*-like odorant response profile, consistent with the presence of *Or85a* in its genome. Second, *D. biarmipes Or67a1* and *Or67a2* responses closely resemble those of their *D. suzukii* orthologs. These receptors are likely expressed in ab10A neurons, as in *D. suzukii*, given that their responses more closely match ab10A than ab2B. Third, *D. biarmipes Or67a5* exhibits robust responses to ethyl benzoate and methyl benzoate, which are the strongest activators of both the ancestral *Or67a* in *D. melanogaster*^27^ and ab10A neurons in *D. biarmipes*. Fourth, although most responses of *D. biarmipes Or67a3* resemble those of *DsuzOr67a3*, ab2B responses in *D. biarmipes* are entirely different. Some responses of *Or67a3* instead resemble those of ab10A, suggesting that its activity may be masked by co-expression with other *or67a* receptors in ab10A.

These differences suggest that the recruitment of *Or67a4* and *Or67a3* into the ab2B neurons is a unique feature to *D. suzukii*, likely emerging after the genomic loss of *Or85a* and the gain of *Or67a4*. As such, *D. biarmipes* represents an intermediate evolutionary state: while it shows early signs of olfactory adaptation in ab3A and ab10A neurons, it does not exhibit β-cyclocitral sensitivity in ab3A neurons, which in *D. suzukii* arises from the co-expression of a second receptor alongside *Or22a*, nor does it show evidence for receptor reassignment in ab2B.

In summary, by co-expressing a second OR with *Or22a* in ab3A and partitioning *Or67a* paralogs between ab10A and ab2B, *D. suzukii* expands its odorant tuning without creating new neuronal classes or rewiring central circuits. This illustrates how gene loss, duplication, functional divergence, and receptor redeployment can reprogram olfactory function to meet the demands of novel ecological niches.

### Limitations of the study

Although we elucidated that *D. suzukii* ab3A neurons express an additional receptor enabling detection of β-cyclocitral, the identity of this receptor remains unknown. Additionally, while our data indicate that *Or67a1* and *Or67a2* are co-expressed in ab10A neurons and that *Or67a4* has replaced the ancestral *Or85a* in ab2B neurons, *Or67a3* is also a strong candidate for ab2B expression, as its ectopic expression produces responses resembling those of native ab2B neurons, and no other ORN responses resemble those of Or67a3. Notably, *Or67a3* lies immediately downstream of *Or67a4*, separated by fewer than 500 base pairs, raising the possibility that our CRISPR-mediated knockout of *Or67a4* may have inadvertently affected *Or67a3* expression. Finally, the regulatory mechanisms governing the partitioned expression of *Or67a* paralogs across these distinct neurons in *D. suzukii* remain unresolved.

## Methods

### Drosophila stocks

Flies, 5-7 days old, were reared on corn syrup and soy four culture medium (Archon Scientifc) at 24°C and 50% relative humidity in a 12:12-h light–dark cycle. *D. melanogaster* Canton-S stock was CS-5. *D. suzukii* Connecticut (CT), North Carolina (NC), and California genome (WT3) strains were obtained from Dr. Richard S. Cowles, Dr. Maxwell J. Scott, and Dr. Joanna C. Chiu, respectively. *D. biarmipes* (14,023– 0361.04) stock was obtained from the Drosophila Species Stock Center. *D. suzukii Orco^3^* mutant line was obtained from Dr. Benjamin Prud’homme. The *D. melanogaster* empty neuron system (*Or22ab^Gal4^*) was obtained from Dr. John R. Carlson.

### Odorants

Odorants of the highest available purity were obtained from Millipore Sigma, TCI America, or Thermo Scientific Chemicals and stored as recommended. A complete list of odorants is provided in Table S1. Odorants were dissolved in hexane (Millipore Sigma, Catalog #139386). Odor presentation was performed by inserting a Pasteur pipette tip into a hole in a 20 cm delivery tube, which directed a continuous airstream over the labium. The airflow through the pipette was controlled using a Stimulus Air Controller CS-55 V2 (Syntech), delivering a 0.5 s pulse. The wind speed at the position of the mounted animal, after adding the odorized stimulus to the continuous airflow, was approximately 50 cm s⁻¹. As an odor source, a cellulose filter disc (∼1 cm diameter) soaked with 10 µl of diluted odorant was placed inside a disposable borosilicate glass Pasteur pipette (2 ml capacity, Fisher Scientific GSA). Odorants were presented sequentially, with at least 60 s between presentations. For dose–response analyses, odorants were delivered in increasing concentrations in logarithmic steps.

### Electrophysiology

A single fly was placed in a 200-μL plastic pipette tip with its head directed towards the narrower end to allow only the antennae to protrude. The pipette tip was then securely attached to a glass microscope slide. The antenna was gently stabilized on a cover slip using a glass capillary. The slide was then placed under a light microscope (BX51WI, Olympus, Tokyo, Japan) equipped with a 50 × objective (LMPLFLN 50X, Olympus) and 10 × eyepieces. A reference tungsten electrode (catalog no. 716000, A-M Systems), electrolytically sharpened to 1 μm tip diameter by dipping it repeatedly in a 10% KNO3 solution, was inserted into the eye. The recording tungsten electrode, identical to the reference electrode, was inserted gently into the base of a coeloconic sensillum. Signals were amplified (10 ×; Syntech Universal AC/DC Probe; http://www.syntech.nl), sampled (10,667 samples s^−1^), and filtered (100–3000 Hz with 50/60 Hz suppression) via a USB-IDAC connection (Syntech) to a computer. Action potentials were extracted using Syntech AutoSpike 32 software. Responses as the increase (or decrease) in the action potential frequency (spikes/s) were calculated by subtracting the number of action potentials during the 0.5 s preceding the odor stimulation from the number of action potentials during the 0.5 s of odor stimulation. We then subtracted the response to hexane from the response to the odorant, and the differences are what we reported.

### Continuous airflow

A 20 cm glass tube with an inner diameter of approximately 4 mm supplied activated charcoal-filtered humidified air to the labium. Air, continuous and pulsed, was generated by a Stimulus Air Controller CS-55 V2 from Syntech (https://www.ockenfels-syntech.com/products/stimulus-controllers/). The end of the tube was placed about 1 cm from the preparation, and the continuous airflow near the mounted animal had a speed of approximately 40 cm s−1.

### Generating UAS-DsuzOr lines

Five *D. suzukii* WT3 females were collected, and whole animals were immediately homogenized for RNA extraction using TRIzol reagent (Invitrogen). The extracted RNA was treated with TURBO DNase (Invitrogen) to remove residual genomic DNA. 1 μg of purified RNA was used for cDNA synthesis with oligo(dT) primers using SuperScript IV Reverse Transcriptase (Invitrogen) for the generation of PCR template for subsequent amplifications of *DsuzOr* genes. PCR reactions were conducted using Phusion High-Fidelity DNA Polymerase (Thermo Scientific) under standard thermal cycling conditions in a Bio-Rad thermocycler. PCR products were separated on 1.0% agarose gel, and DNA fragments of the expected sizes were excised and purified according to the manufacturer’s instructions (Wizard SV Gel and PCR Cleanup System, Promega).

*DsuzOrs* were cloned into the pJFRC81-10XUAS-IVS-Syn21-GFP-p10 vector (pJFRC81, Addgene plasmid #36432). Forward and reverse PCR primers of *DsuzOrs* were designed with 20 bp overhangs complementary to the regions flanking the GFP coding sequence in pJFRC81. The pJFRC81 vector was linearized by PCR using primers flanking the GFP sequence, with effective removal of the GFP coding region. Gibson assembly (NEB) was then used to ligate the amplified *DsuzOr* fragments into the linearized vector. The resulting constructs were transformed into bacteria and sequence-verified. Verified plasmids were injected into embryos of BDSC Stock #8622 by BestGene Inc. for PhiC31-mediated site-specific integration. Primers are provided in Table S2.

### CRISPR–Cas9-mediated genome engineering

Single guide RNAs (sgRNAs) targeting *DsuzOrs* were designed using the online CHOPCHOP tool (https://chopchop.cbu.uib.no/). sgRNAs were synthesized through *in vitro* transcription using the GeneArt Precision gRNA Synthesis Kit (Thermo Fisher Scientific). The sgRNA was mixed with Cas9 protein (TrueCut, Invitrogen) at a final concentration of 200 ng/μl and 500ng/μl, respectively, in a 10 μl reaction with 1 μl of injection buffer (300 mM sodium phosphate and 1 mM potassium chloride) and an additional 1 μl of 2M potassium chloride. The sgRNA/Cas9 mixture was incubated at 37°C for 10 min in a thermocycler to promote ribonucleoprotein (RNP) complex formation. The resulting RNP solution was passed through a Millipore Ultrafree-MC filter (0.45 μM pore size) and kept on ice until ready to use.

The microinjection of RNPs into *Drosophila* embryos was performed using a FemtoJet 4i microinjector (Eppendorf). Injection needles were prepared from capillary tubes (World Precision Instrument, 1B100F-4) using a micropipette puller (Model P-1000, Sutter Instrument). One day before microinjection, approximately 300 flies (3-5 days old) were transferred to an egg-laying chamber (Flystuff Cat #59-100) containing a Petri dish filled with 10 ml of egg-laying substrate composed of 25g agar and 200ml molasses dissolved in 500 ml of water. To diminish the older eggs retained in the females’ ovaries, the egg-laying Petri dish was replaced every 20 min for at least 5 times before injection. Only embryos laid within 20 min were used for injection, and the entire injection process was completed within 1h.

G0 females that emerged from injected embryos (G0) were crossed to wild-type flies to generate G1 heterozygotes. G1 individuals with potential CRISPR-induced mutations were crossed again to wild-type flies to produce G2 heterozygotes. G1 flies were sacrificed for genotyping to confirm the mutation types. G2 heterozygotes were self-crossed to obtain homozygous mutants and establish stable mutant lines. Mutants were identified using a PCR-based screening strategy. Primers for sgRNA synthesis and mutant screening are listed in Table S3.

### Structural prediction and molecular docking

The 3D homotetrameric structures of *DsuzOr22a^WT(D201A)^* and *DsuzOr22a^A201D^* were generated with AlphaFold2^50^. The highest-confidence predictions (*model_0*) were selected for docking. The ligand, isobutyl acetate (SDF file), was obtained from PubChem (https://pubchem.ncbi.nlm.nih.gov/) and converted to mol2 format with OpenBabel (v3.1.1)^51^. Structural preparation of both the ligand and receptor subunits was performed in PyMol (v3.1.4), and the resulting files were converted to PDBQT format with AutoDockTools (v1.5.7). Molecular docking was carried out using AutoDock Vina (v1.2.7) with a cubic grid box (20 Å x 20 Å x 20 Å) centered on the transmembrane domains. Receptor–ligand interactions were analyzed with the Protein-Ligand Interaction Profiler (v2.4.0) and visualized in PyMol (v3.1.4).

### Maximum-likelihood phylogenetic trees

To construct maximum-likelihood phylogenetic trees of the Or67a subfamily, we retrieved the amino acid sequences of D. suzukii and D. biarmipes Or67a paralogs, as well as D. melanogaster Or67a, from NCBI. Sequences were aligned using ClustalW in MEGA version 11 with default parameters. We then used MEGA’s “Find Best Protein Models (ML)” to select the best-fit substitution model based on the lowest Bayesian Information Criterion (BIC) score. This model was applied to generate maximum-likelihood trees with 1,000 bootstrap replicates to assess branch support.

### Quantification and statistical analysis

The heatmaps were generated in PAST. Euclidean distances were calculated in R (version 4.2.3) using the ‘moments’ package, and other statistical tests were performed in GraphPad Prism (version 10.0.1 [316]).

## Acknowledgements

This work was supported by NIH grant K01 DC020145 to H.K.M.D.

## Contributions

H.K.M.D. conceived the study. H.K.M.D. and Q.X. designed the experiments. H.K.M.D. and Q.X. carried out the experiments. H.K.M.D. and Q.X. analysed the data. H.K.M.D. and Q.X. prepared the original draft of the manuscript. H.K.M.D. and Q.X. read and approved the final version of the manuscript.

## Supplementary Tables and Figures

**Table 1.**
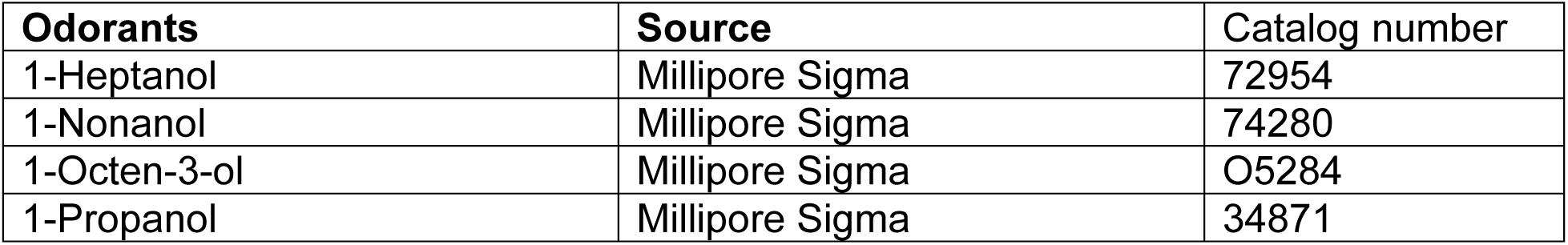
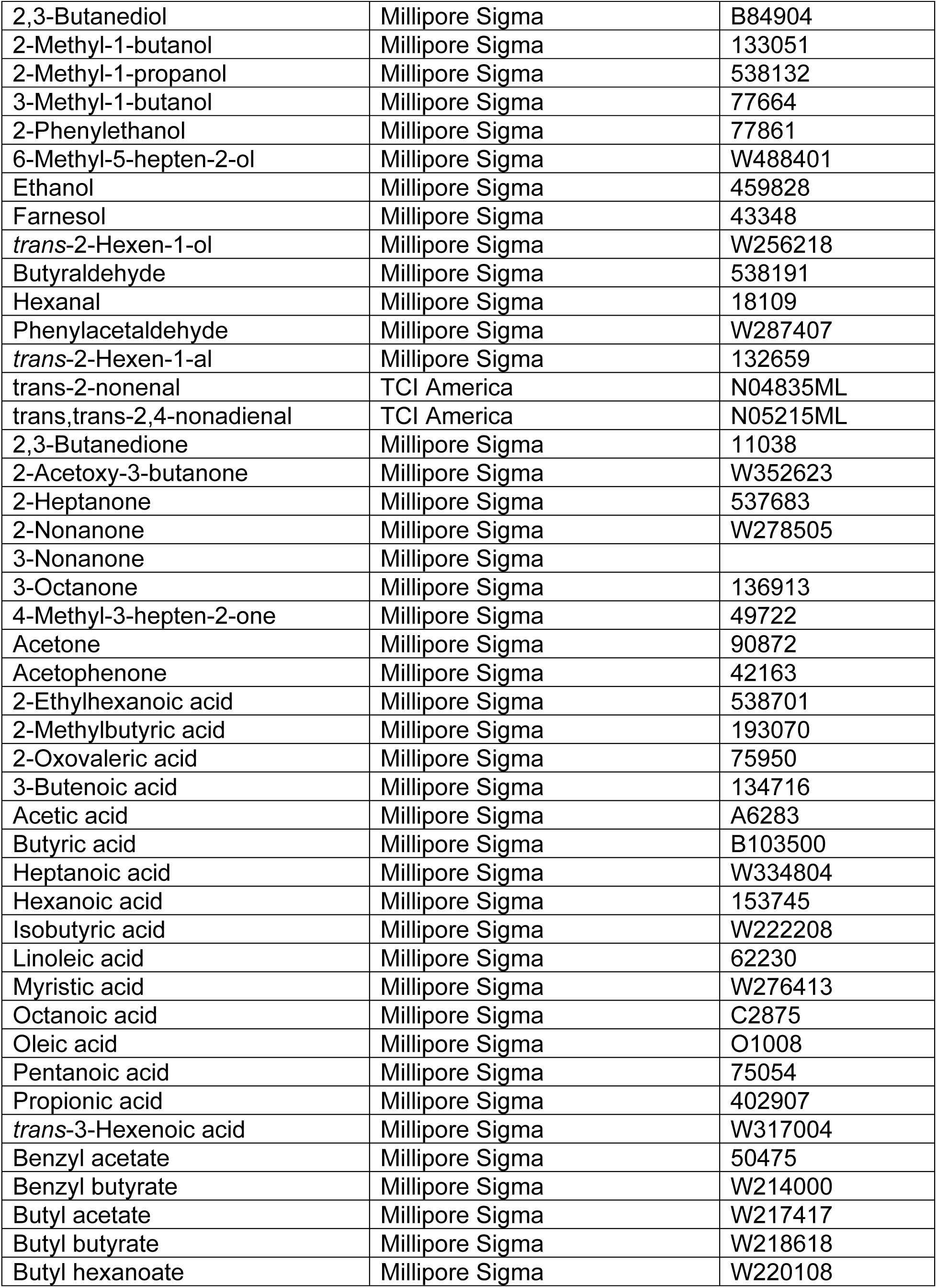
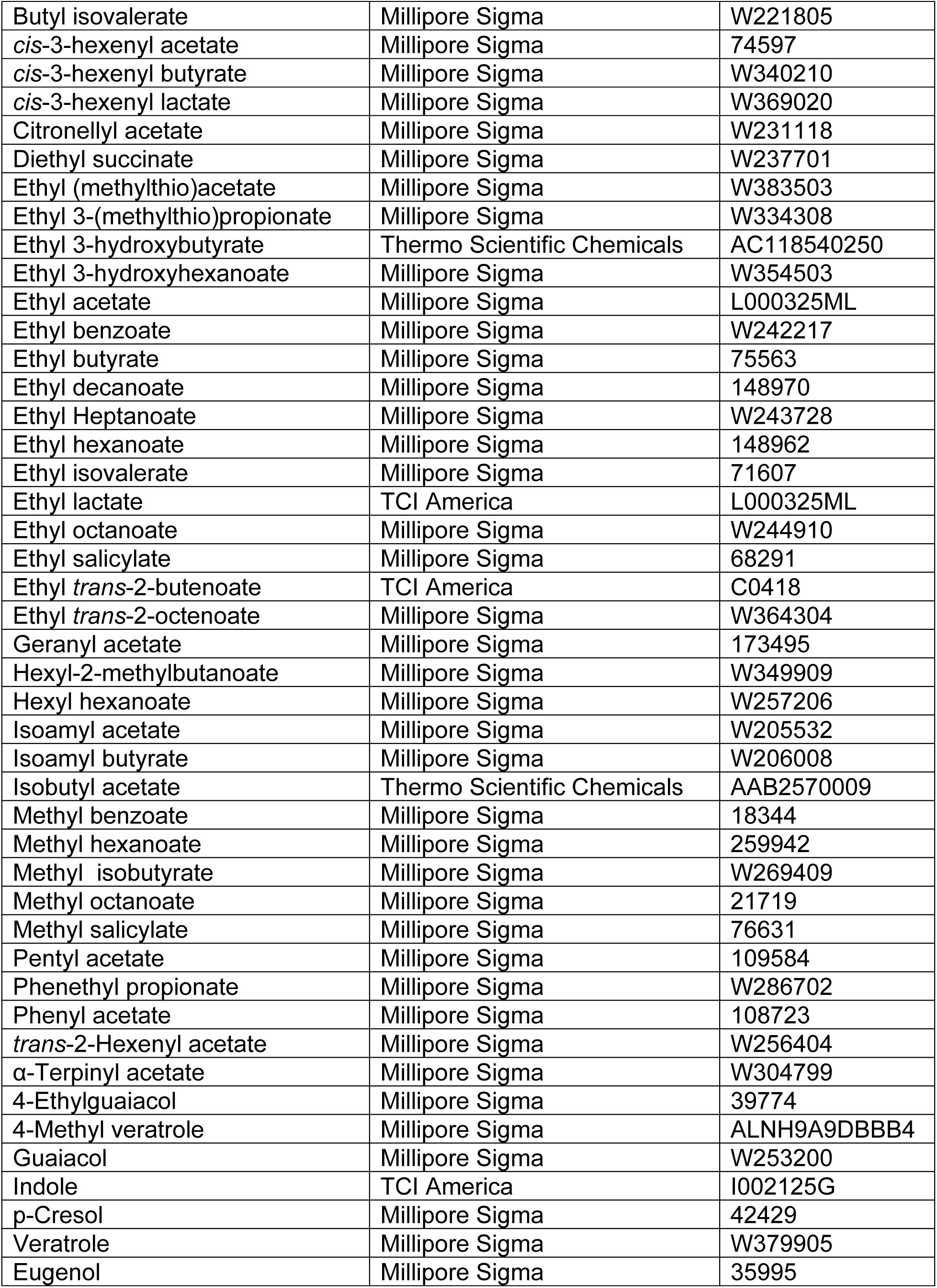
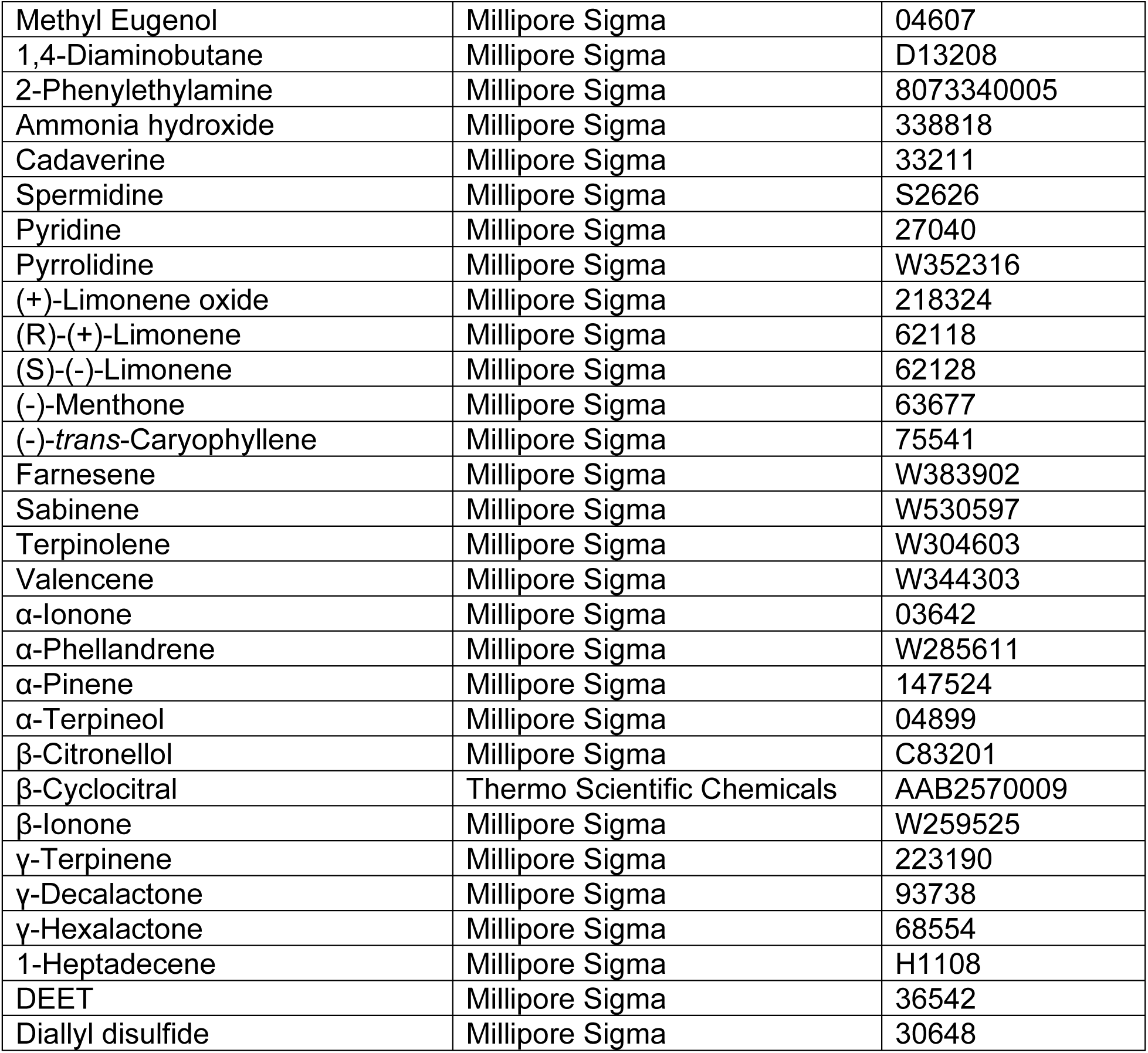
List of odorants.

| Odorants | Source | Catalog number |
| --- | --- | --- |
| 1-Heptanol | Millipore Sigma | 72954 |
| 1-Nonanol | Millipore Sigma | 74280 |
| 1-Octen-3-ol | Millipore Sigma | O5284 |
| 1-Propanol | Millipore Sigma | 34871 |
| 2,3-Butanediol | Millipore Sigma | B84904 |
| 2-Methyl-1-butanol | Millipore Sigma | 133051 |
| 2-Methyl-1-propanol | Millipore Sigma | 538132 |
| 3-Methyl-1-butanol | Millipore Sigma | 77664 |
| 2-Phenylethanol | Millipore Sigma | 77861 |
| 6-Methyl-5-hepten-2-ol | Millipore Sigma | W488401 |
| Ethanol | Millipore Sigma | 459828 |
| Farnesol | Millipore Sigma | 43348 |
| *trans*-2-Hexen-1-ol | Millipore Sigma | W256218 |
| Butyraldehyde | Millipore Sigma | 538191 |
| Hexanal | Millipore Sigma | 18109 |
| Phenylacetaldehyde | Millipore Sigma | W287407 |
| *trans*-2-Hexen-1-al | Millipore Sigma | 132659 |
| *trans*-2-nonenal | TCI America | N04835ML |
| *trans,trans*-2,4-nonadienal | TCI America | N05215ML |
| 2,3-Butanedione | Millipore Sigma | 11038 |
| 2-Acetoxy-3-butanone | Millipore Sigma | W352623 |
| 2-Heptanone | Millipore Sigma | 537683 |
| 2-Nonanone | Millipore Sigma | W278505 |
| 3-Nonanone | Millipore Sigma |  |
| 3-Octanone | Millipore Sigma | 136913 |
| 4-Methyl-3-hepten-2-one | Millipore Sigma | 49722 |
| Acetone | Millipore Sigma | 90872 |
| Acetophenone | Millipore Sigma | 42163 |
| 2-Ethylhexanoic acid | Millipore Sigma | 538701 |
| 2-Methylbutyric acid | Millipore Sigma | 193070 |
| 2-Oxovaleric acid | Millipore Sigma | 75950 |
| 3-Butenoic acid | Millipore Sigma | 134716 |
| Acetic acid | Millipore Sigma | A6283 |
| Butyric acid | Millipore Sigma | B103500 |
| Heptanoic acid | Millipore Sigma | W334804 |
| Hexanoic acid | Millipore Sigma | 153745 |
| Isobutyric acid | Millipore Sigma | W222208 |
| Linoleic acid | Millipore Sigma | 62230 |
| Myristic acid | Millipore Sigma | W276413 |
| Octanoic acid | Millipore Sigma | C2875 |
| Oleic acid | Millipore Sigma | O1008 |
| Pentanoic acid | Millipore Sigma | 75054 |
| Propionic acid | Millipore Sigma | 402907 |
| *trans*-3-Hexenoic acid | Millipore Sigma | W317004 |
| Benzyl acetate | Millipore Sigma | 50475 |
| Benzyl butyrate | Millipore Sigma | W214000 |
| Butyl acetate | Millipore Sigma | W217417 |
| Butyl butyrate | Millipore Sigma | W218618 |
| Butyl hexanoate | Millipore Sigma | W220108 |
| Butyl isovalerate | Millipore Sigma | W221805 |
| *cis*-3-hexenyl acetate | Millipore Sigma | 74597 |
| *cis*-3-hexenyl butyrate | Millipore Sigma | W340210 |
| *cis*-3-hexenyl lactate | Millipore Sigma | W369020 |
| Citronellyl acetate | Millipore Sigma | W231118 |
| Diethyl succinate | Millipore Sigma | W237701 |
| Ethyl (methylthio)acetate | Millipore Sigma | W383503 |
| Ethyl 3-(methylthio)propionate | Millipore Sigma | W334308 |
| Ethyl 3-hydroxybutyrate | Thermo Scientific Chemicals | AC118540250 |
| Ethyl 3-hydroxyhexanoate | Millipore Sigma | W354503 |
| Ethyl acetate | Millipore Sigma | L000325ML |
| Ethyl benzoate | Millipore Sigma | W242217 |
| Ethyl butyrate | Millipore Sigma | 75563 |
| Ethyl decanoate | Millipore Sigma | 148970 |
| Ethyl Heptanoate | Millipore Sigma | W243728 |
| Ethyl hexanoate | Millipore Sigma | 148962 |
| Ethyl isovalerate | Millipore Sigma | 71607 |
| Ethyl lactate | TCI America | L000325ML |
| Ethyl octanoate | Millipore Sigma | W244910 |
| Ethyl salicylate | Millipore Sigma | 68291 |
| Ethyl *trans*-2-butenate | TCI America | C0418 |
| Ethyl *trans*-2-octenoate | Millipore Sigma | W364304 |
| Geranyl acetate | Millipore Sigma | 173495 |
| Hexyl-2-methylbutanoate | Millipore Sigma | W349909 |
| Hexyl hexanoate | Millipore Sigma | W257206 |
| Isoamyl acetate | Millipore Sigma | W205532 |
| Isoamyl butyrate | Millipore Sigma | W206008 |
| Isobutyl acetate | Thermo Scientific Chemicals | AAB2570009 |
| Methyl benzoate | Millipore Sigma | 18344 |
| Methyl hexanoate | Millipore Sigma | 259942 |
| Methyl isobutyrate | Millipore Sigma | W269409 |
| Methyl octanoate | Millipore Sigma | 21719 |
| Methyl salicylate | Millipore Sigma | 76631 |
| Pentyl acetate | Millipore Sigma | 109584 |
| Phenethyl propionate | Millipore Sigma | W286702 |
| Phenyl acetate | Millipore Sigma | 108723 |
| *trans*-2-Hexenyl acetate | Millipore Sigma | W256404 |
| α-Terpinyl acetate | Millipore Sigma | W304799 |
| 4-Ethylguaiaicol | Millipore Sigma | 39774 |
| 4-Methyl veratrole | Millipore Sigma | ALNH9A9DBBB4 |
| Guaiaicol | Millipore Sigma | W253200 |
| Indole | TCI America | I002125G |
| p-Cresol | Millipore Sigma | 42429 |
| Veratrole | Millipore Sigma | W379905 |
| Eugenol | Millipore Sigma | 35995 |
| Methyl Eugenol | Millipore Sigma | 04607 |
| 1,4-Diaminobutane | Millipore Sigma | D13208 |
| 2-Phenylethylamine | Millipore Sigma | 8073340005 |
| Ammonia hydroxide | Millipore Sigma | 338818 |
| Cadaverine | Millipore Sigma | 33211 |
| Spermidine | Millipore Sigma | S2626 |
| Pyridine | Millipore Sigma | 27040 |
| Pyrrolidine | Millipore Sigma | W352316 |
| (+)-Limonene oxide | Millipore Sigma | 218324 |
| (R)-(+)-Limonene | Millipore Sigma | 62118 |
| (S)-(-)-Limonene | Millipore Sigma | 62128 |
| (-)-Menthone | Millipore Sigma | 63677 |
| (-)-*trans*-Caryophyllene | Millipore Sigma | 75541 |
| Farnesene | Millipore Sigma | W383902 |
| Sabinene | Millipore Sigma | W530597 |
| Terpinolene | Millipore Sigma | W304603 |
| Valencene | Millipore Sigma | W344303 |
| α-Ionone | Millipore Sigma | 03642 |
| α-Phellandrene | Millipore Sigma | W285611 |
| α-Pinene | Millipore Sigma | 147524 |
| α-Terpineol | Millipore Sigma | 04899 |
| β-Citronellol | Millipore Sigma | C83201 |
| β-Cyclocitral | Thermo Scientific Chemicals | AAB2570009 |
| β-Ionone | Millipore Sigma | W259525 |
| γ-Terpinene | Millipore Sigma | 223190 |
| γ-Decalactone | Millipore Sigma | 93738 |
| γ-Hexalactone | Millipore Sigma | 68554 |
| 1-Heptadecene | Millipore Sigma | H1108 |
| DEET | Millipore Sigma | 36542 |
| Diallyl disulfide | Millipore Sigma | 30648 |

**Table 2.**
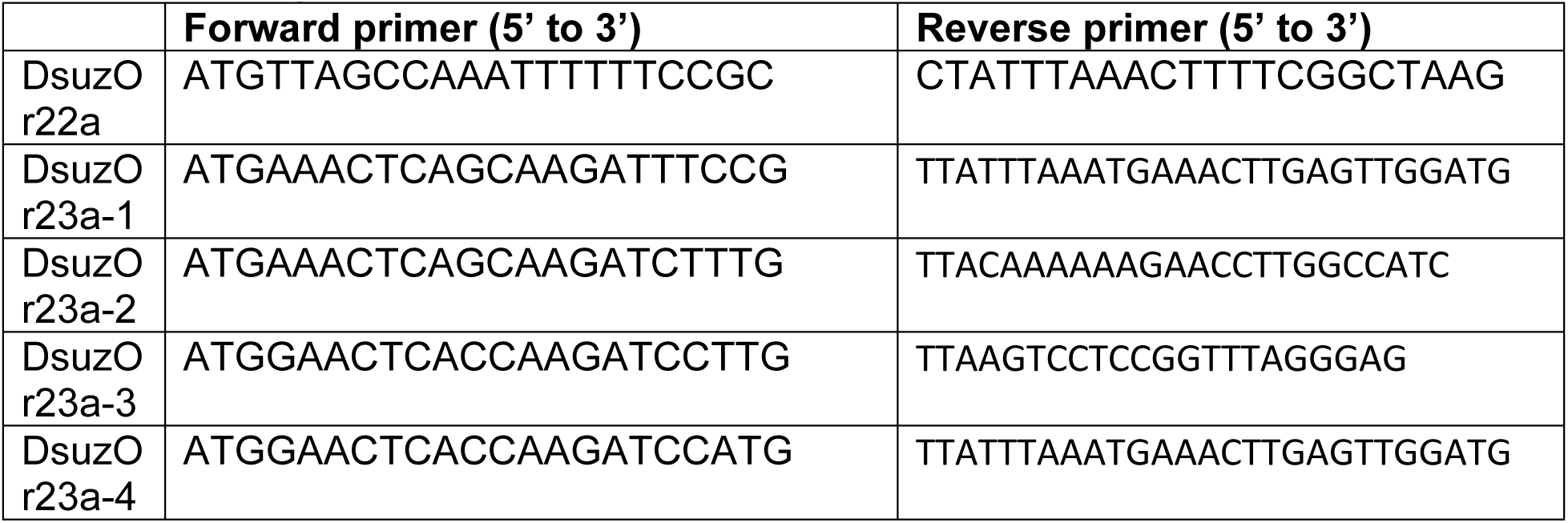
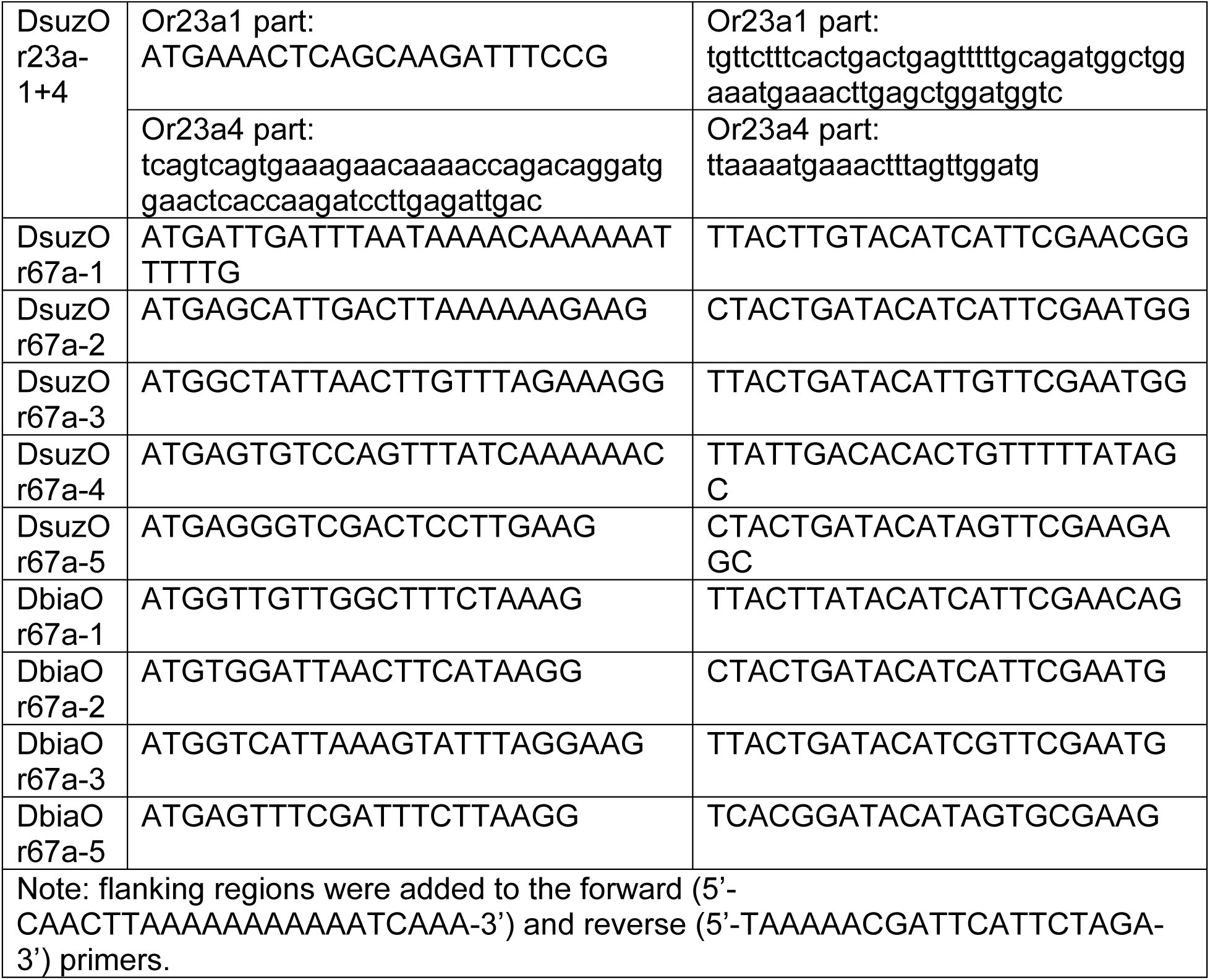
List of primers for UAS constructs.

**Table 3.**
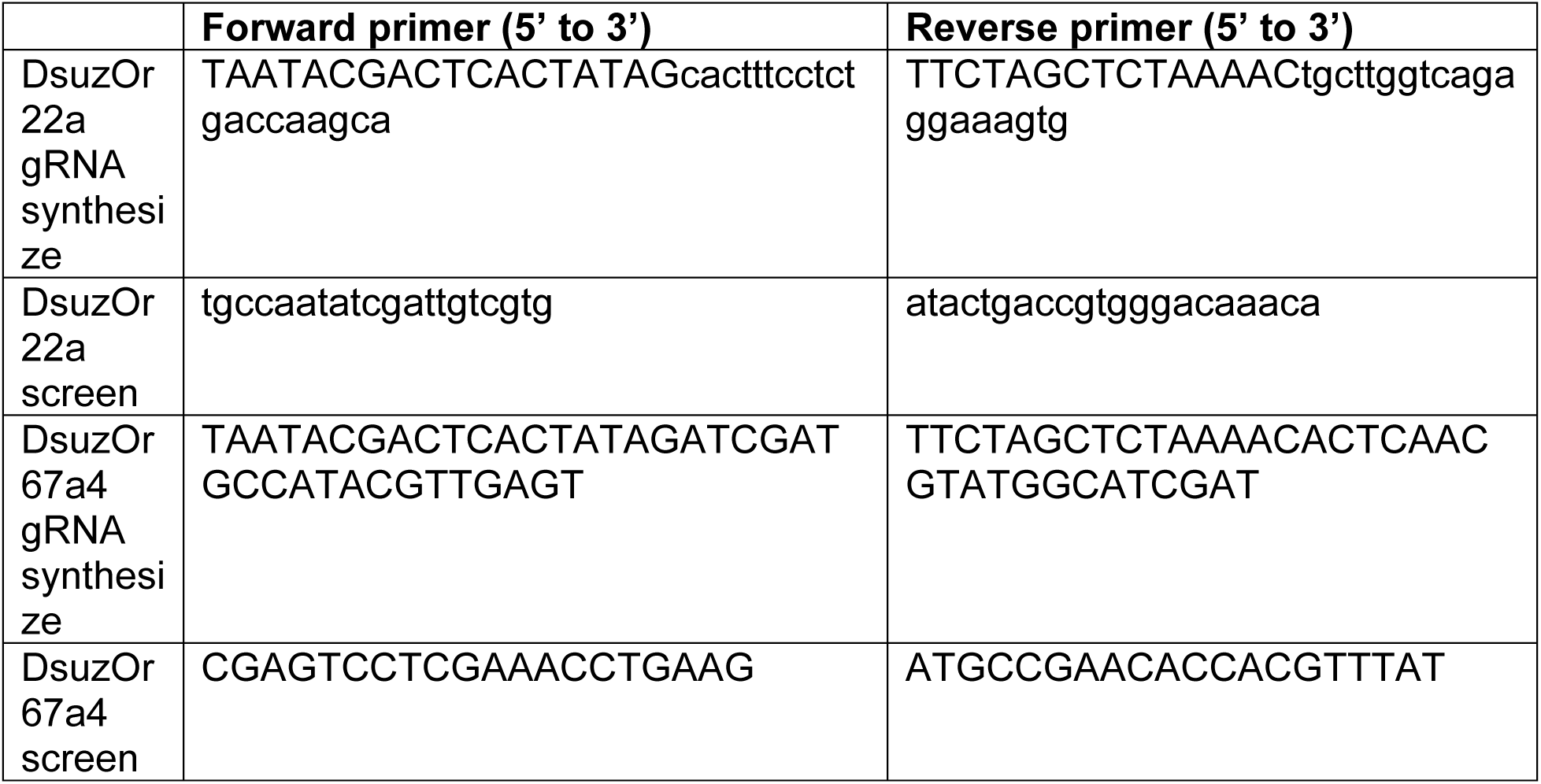
Primers for CRISPR.

**Figure S1.**
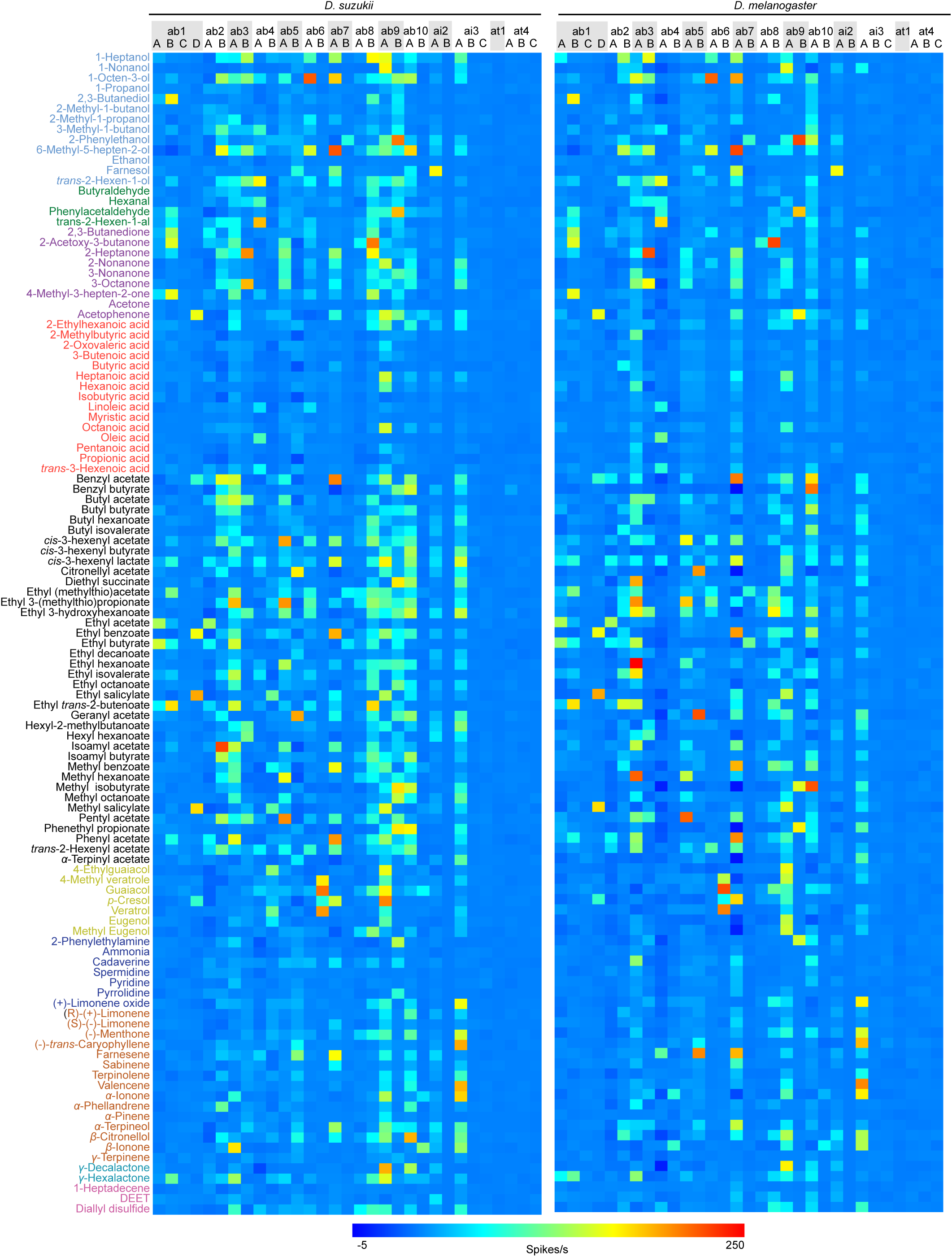
Responses of ORNs housed in D. suzukii and *D. melanogaster* antennal Basiconic, and intermediate, and trichoid sensilla. (A,B) Heatmaps of responses of ORNs housed in *D. suzukii* (A) and *D. melanogaster* (C) antennal basiconic, intermediate, and trichoid sensilla to 113 odorants. Each odorant was diluted in hexane and tested at a 10^-2^ dilution. Odorants are color coded by functional group.

**Figure S2.**
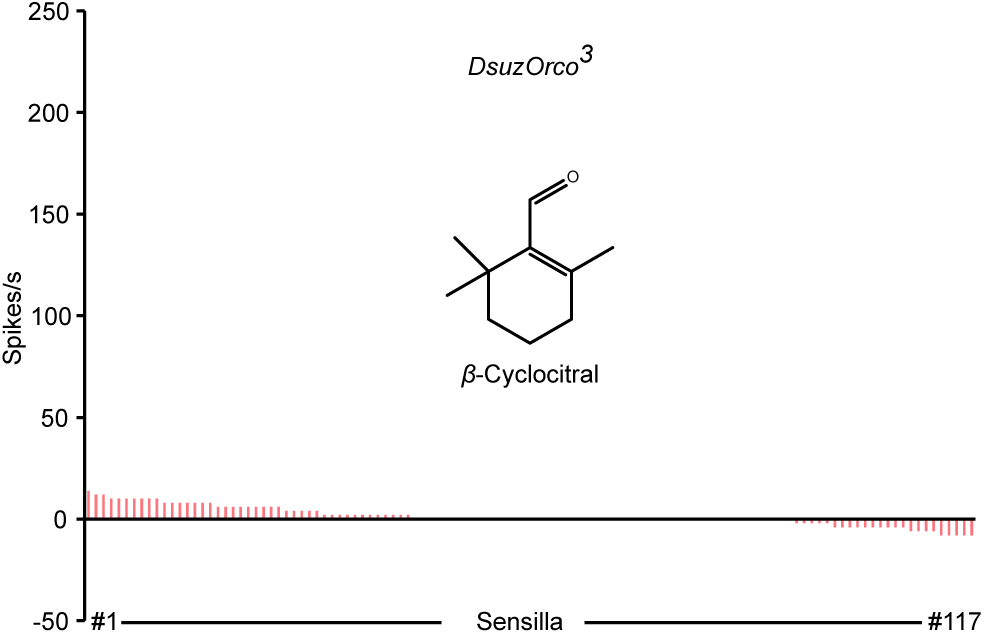
Responses of 117 individual large basiconic sensilla in *DsuzOrco^3^* to β-cyclocitral.

**Figure S3.**
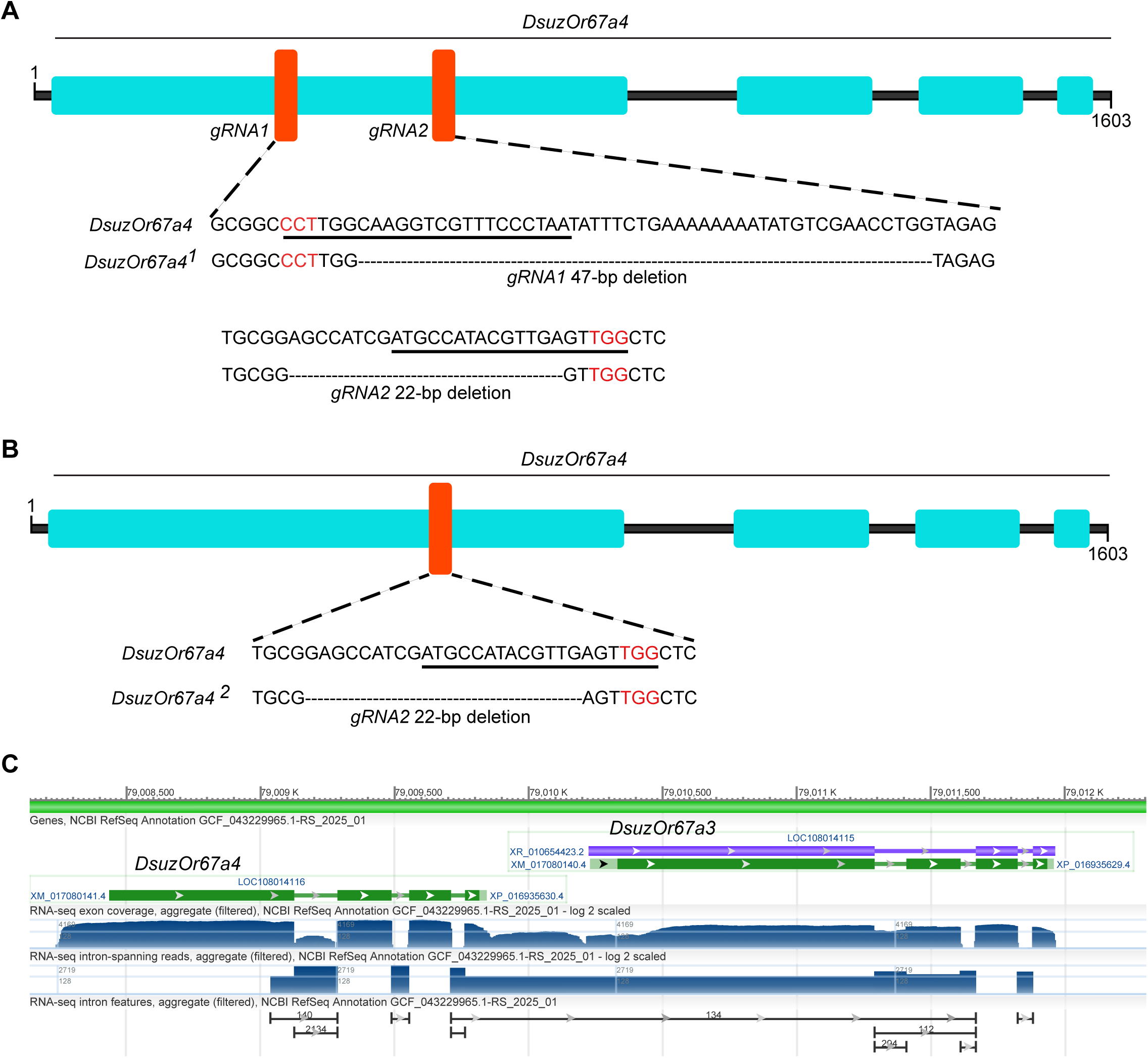
Generation of *DsuzOr67a4* mutant alleles. (A,B) Schematics illustrating the generation of *DsuzOr67a4^1^*(A) *and DsuzOr67a4^2^* (B) mutants using CRISPR/Cas9-mediated gene disruption. Nucleotides in red represent the PAM site, underlined nucleotides indicate the gRNA sequence, and dashed lines denote deleted nucleotides. (D) RNA sequencing data for *DsuzOr67a4* and *DsuzOr67a3*. Green boxes represent exons, and green lines represent introns.

## References

1 Benton, R. *Drosophila* olfaction: past, present and future. Proc Biol Sci 289, 20222054, doi:10.1098/rspb.2022.2054 (2022).

2 Hansson, B. S. & Stensmyr, M. C. Evolution of insect olfaction. Neuron 72, 698–711, doi:10.1016/j.neuron.2011.11.003 (2011).

3 Joseph, R. M. & Carlson, J. R. *Drosophila* Chemoreceptors: A Molecular Interface Between the Chemical World and the Brain. Trends Genet 31, 683–695, doi:10.1016/j.tig.2015.09.005 (2015).

4 Montell, C. Drosophila sensory receptors-a set of molecular Swiss Army Knives. Genetics 217, 1–34, doi:10.1093/genetics/iyaa011 (2021).

5 Andersson, M. N., Löfstedt, C. & Newcomb, R. D. Insect olfaction and the evolution of receptor tuning. Frontiers in Ecology and Evolution Volume 3 - 2015, doi:10.3389/fevo.2015.00053 (2015).

6 Auer, T. O., Shahandeh, M. P. & Benton, R. *Drosophila sechellia*: A Genetic Model for Behavioral Evolution and Neuroecology. Annu Rev Genet 55, 527–554, doi:10.1146/annurev-genet-071719-020719 (2021).

7 Keesey, I. W. Sensory neuroecology and multimodal evolution across the genus Drosophila. Frontiers in Ecology and Evolution Volume 10 - 2022, doi:10.3389/fevo.2022.932344 (2022).

8 Nozawa, M. & Nei, M. Evolutionary dynamics of olfactory receptor genes in *Drosophila* species. Proc Natl Acad Sci U S A 104, 7122–7127, doi:10.1073/pnas.0702133104 (2007).

9 Peláez, J. N. et al. Evolution of chemosensory and detoxification gene families across herbivorous Drosophilidae. G3 Genes|Genomes|Genetics 13, doi:10.1093/g3journal/jkad133 (2023).

10 Sánchez-Gracia, A., Vieira, F. G. & Rozas, J. Molecular evolution of the major chemosensory gene families in insects. Heredity 103, 208–216, doi:10.1038/hdy.2009.55 (2009).

11 Eyun, S. I. et al. Evolutionary history of chemosensory-related gene families across the Arthropoda. Mol Biol Evol 34, 1838–1862, doi:10.1093/molbev/msx147 (2017).

12 McBride, C. S., Arguello, J. R. & O’Meara, B. C. Five Drosophila genomes reveal nonneutral evolution and the signature of host specialization in the chemoreceptor superfamily. Genetics 177, 1395–1416, doi:10.1534/genetics.107.078683 (2007).

13 Dweck, H. K., Talross, G. J., Wang, W. & Carlson, J. R. Evolutionary shifts in taste coding in the fruit pest *Drosophila suzukii*. Elife 10, doi:10.7554/eLife.64317 (2021).

14 Karageorgi, M. et al. Evolution of Multiple Sensory Systems Drives Novel Egg-Laying Behavior in the Fruit Pest *Drosophila suzukii*. Curr Biol 27, 847–853, doi:10.1016/j.cub.2017.01.055 (2017).

15 Yeh, D. A., Drummond, F. A., Gómez, M. I. & Fan, X. The economic impacts and management of spotted wing *Drosophila* (*Drosophila Suzukii*): The case of wild blueberries in Maine. Journal of Economic Entomology 113, 1262–1269, doi:10.1093/jee/toz360 (2020).

16 Auer, T. O., Álvarez-Ocaña, R., Cruchet, S., Benton, R. & Arguello, J. R. Copy number changes in co-expressed odorant receptor genes enable selection for sensory differences in *drosophilid* species. Nat Ecol Evol 6, 1343–1353, doi:10.1038/s41559-022-01830-y (2022).

17 Goldman-Huertas, B. et al. Evolution of herbivory in *Drosophilidae* linked to loss of behaviors, antennal responses, odorant receptors, and ancestral diet. Proc Natl Acad Sci U S A 112, 3026–3031, doi:10.1073/pnas.1424656112 (2015).

18 Matsunaga, T. et al. Evolution of Olfactory Receptors Tuned to Mustard Oils in Herbivorous Drosophilidae. Mol Biol Evol 39, doi:10.1093/molbev/msab362 (2022).

19. 19 Matsunaga, T., et al. Odorant receptors mediating avoidance of toxic mustard oils in *Drosophila melanogaster* are expanded in herbivorous relatives. bioRxiv, doi:10.1101/2024.10.08.617316 (2025).

20 Shaw, K. H., Johnson, T. K., Anderson, A., de Bruyne, M. & Warr, C. G. Molecular and Functional Evolution at the Odorant Receptor Or22 Locus in *Drosophila melanogaster*. Molecular Biology and Evolution 36, 919–929, doi:10.1093/molbev/msz018 (2019).

21 Durkin, S. M. et al. Behavioral and Genomic Sensory Adaptations Underlying the Pest Activity of *Drosophila suzukii*. Mol Biol Evol 38, 2532–2546, doi:10.1093/molbev/msab048 (2021).

22 Hickner, P. V. et al. The making of a pest: Insights from the evolution of chemosensory receptor families in a pestiferous and invasive fly, *Drosophila suzukii*. BMC Genomics 17, 648, doi:10.1186/s12864-016-2983-9 (2016).

23 Ramasamy, S. et al. The Evolution of olfactory gene families in *Drosophila* and the genomic basis of chemical-Ecological adaptation in *Drosophila suzukii*. Genome Biol Evol 8, 2297–2311, doi:10.1093/gbe/evw160 (2016).

24 Walker, W. B., 3rd et al. Comparative transcriptomic assessment of the chemosensory receptor repertoire of *Drosophila suzukii* adult and larval olfactory organs. Comp Biochem Physiol Part D Genomics Proteomics 45, 101049, doi:10.1016/j.cbd.2022.101049 (2023).

25 Benton, R. et al. An integrated anatomical, functional and evolutionary view of the *Drosophila* olfactory system. EMBO reports 0, 1–22, 10.1038/s44319-025-00476-8 (2025).

26 Dweck, H. K. M. et al. The Olfactory Logic behind Fruit Odor Preferences in Larval and Adult *Drosophila*. Cell Rep 23, 2524–2531, doi:10.1016/j.celrep.2018.04.085 (2018).

27 Hallem, E. A. & Carlson, J. R. Coding of odors by a receptor repertoire. Cell 125, 143–160, doi:10.1016/j.cell.2006.01.050 (2006).

28 Hickner, P. V. et al. The making of a pest: Insights from the evolution of chemosensory receptor families in a pestiferous and invasive fly, *Drosophila suzukii*. BMC Genomics 17, 648, doi:10.1186/s12864-016-2983-9 (2016).

29 Dweck, H. K. et al. Olfactory channels associated with the *Drosophila* maxillary palp mediate short- and long-range attraction. Elife 5, doi:10.7554/eLife.14925 (2016).

30 Xue, Q., Hasan, K. S., Dweck, O., Ebrahim, S. A. M. & Dweck, H. K. M. Functional characterization and evolution of olfactory responses in coeloconic sensilla of the global fruit pest *Drosophila suzukii*. BMC Biol 23, 50, doi:10.1186/s12915-025-02151-9 (2025).

31 Keesey, I. W., Knaden, M. & Hansson, B. S. Olfactory specialization in *Drosophila suzukii* supports an ecological shift in host preference from rotten to fresh fruit. J Chem Ecol 41, 121–128, doi:10.1007/s10886-015-0544-3 (2015).

32 Keesey, I. W. et al. Functional olfactory evolution in *Drosophila* suzukii and the subgenus Sophophora. iScience 25, 104212, doi:10.1016/j.isci.2022.104212 (2022).

33 Depetris-Chauvin, A. et al. Evolution at multiple processing levels underlies odor-guided behavior in the genus *Drosophila*. Curr Biol 33, 4771–4785 e4777, doi:10.1016/j.cub.2023.09.039 (2023).

34 Chiu, J. C. et al. Genome of Drosophila suzukii, the spotted wing drosophila. G3 (Bethesda) 3, 2257–2271, doi:10.1534/g3.113.008185 (2013).

35 Ometto, L. et al. Linking genomics and ecology to investigate the complex evolution of an invasive *Drosophila* pest. Genome Biol Evol 5, 745–757, doi:10.1093/gbe/evt034 (2013).

36 Versace, E. et al. Physiological and behavioral responses in *Drosophila melanogaster* to odorants present at different plant maturation stages. Physiol Behav 163, 322–331, doi:10.1016/j.physbeh.2016.05.027 (2016).

37 Cha, D. H., Adams, T., Rogg, H. & Landolt, P. J. Identification and field evaluation of fermentation volatiles from wine and vinegar that mediate attraction of spotted wing *Drosophila, Drosophila suzukii*. J Chem Ecol 38, 1419–1431, doi:10.1007/s10886-012-0196-5 (2012).

38 Chahda, J. S. et al. The molecular and cellular basis of olfactory response to tsetse fly attractants. PLoS Genet 15, e1008005, doi:10.1371/journal.pgen.1008005 (2019).

39 Shaw, K. H., Johnson, T. K., Anderson, A., de Bruyne, M. & Warr, C. G. Molecular and Functional Evolution at the Odorant Receptor *Or22* Locus in *Drosophila melanogaster*. Mol Biol Evol 36, 919–929, doi:10.1093/molbev/msz018 (2019).

40 Butterwick, J. A. et al. Cryo-EM structure of the insect olfactory receptor Orco. Nature 560, 447–452, doi:10.1038/s41586-018-0420-8 (2018).

41 Crava, C. M., Sassù, F., Tait, G., Becher, P. G. & Anfora, G. Functional transcriptome analyses of *Drosophila suzukii* antennae reveal mating-dependent olfaction plasticity in females. Insect Biochem Mol Biol 105, 51–59, doi:10.1016/j.ibmb.2018.12.012 (2019).

42 Dweck, H. K. M. & Carlson, J. R. Molecular Logic and Evolution of Bitter Taste in *Drosophila*. Curr Biol 30, 17–30.e13, doi:10.1016/j.cub.2019.11.005 (2020).

43 Dekker, T., Ibba, I., Siju, K. P., Stensmyr, M. C. & Hansson, B. S. Olfactory shifts parallel superspecialism for toxic fruit in *Drosophila melanogaster* sibling, D. sechellia. Curr Biol 16, 101–109, doi:10.1016/j.cub.2005.11.075 (2006).

44 Stensmyr, M. C., Dekker, T. & Hansson, B. S. Evolution of the olfactory code in the *Drosophila melanogaster* subgroup. Proc Biol Sci 270, 2333–2340, doi:10.1098/rspb.2003.2512 (2003).

45 Auer, T. O. et al. Olfactory receptor and circuit evolution promote host specialization. Nature 579, 402–408, doi:10.1038/s41586-020-2073-7 (2020).

46 Shaw, K. H. et al. Natural variation at the *Drosophila melanogaster Or22* odorant receptor locus is associated with changes in olfactory behaviour. Open Biol 11, 210158, doi:10.1098/rsob.210158 (2021).

47 Mansourian, S. et al. Wild African *Drosophila melanogaster* Are Seasonal Specialists on Marula Fruit. Curr Biol 28, 3960–3968 e3963, doi:10.1016/j.cub.2018.10.033 (2018).

48 Crowley-Gall, A. et al. Olfactory variation among closely related cactophilic *Drosophila* species. J Comp Physiol A Neuroethol Sens Neural Behav Physiol, doi:10.1007/s00359-025-01744-7 (2025).

49 de Bruyne, M., Smart, R., Zammit, E. & Warr, C. G. Functional and molecular evolution of olfactory neurons and receptors for aliphatic esters across the *Drosophila* genus. J Comp Physiol A Neuroethol Sens Neural Behav Physiol 196, 97–109, doi:10.1007/s00359-009-0496-6 (2010).

50 Jumper, J. et al. Highly accurate protein structure prediction with AlphaFold. Nature 596, 583–589, doi:10.1038/s41586-021-03819-2 (2021).

51 O’Boyle, N. M. et al. Open Babel: An open chemical toolbox. J Cheminform 3, 33, doi:10.1186/1758-2946-3-33 (2011).

